# Discovery and Evaluation of Biosynthetic Pathways for the Production of Five Methyl Ethyl Ketone Precursors

**DOI:** 10.1101/209569

**Authors:** Milenko Tokic, Noushin Hadadi, Meric Ataman, Dário Neves, Birgitta E. Ebert, Lars M. Blank, Ljubisa Miskovic, Vassily Hatzimanikatis

## Abstract

The limited supply of fossil fuels and the establishment of new environmental policies shifted research in industry and academia towards sustainable production of the 2^nd^ generation of biofuels, with Methyl Ethyl Ketone (MEK) being one promising fuel candidate. MEK is a commercially valuable petrochemical with an extensive application as a solvent. However, as of today, a sustainable and economically viable production of MEK has not yet been achieved despite several attempts of introducing biosynthetic pathways in industrial microorganisms. We used BNICE.ch as a retrobiosynthesis tool to discover all novel pathways around MEK. Out of 1’325 identified compounds connecting to MEK with one reaction step, we selected 3-oxopentanoate, but-3-en-2-one, but-1-en-2-olate, butylamine, and 2-hydroxy-2-methyl-butanenitrile for further study. We reconstructed 3’679’610 novel biosynthetic pathways towards these 5 compounds. We then embedded these pathways into the genome-scale model of *E. coli*, and a set of 18’622 were found to be most biologically feasible ones based on thermodynamics and their yields. For each novel reaction in the viable pathways, we proposed the most similar KEGG reactions, with their gene and protein sequences, as candidates for either a direct experimental implementation or as a basis for enzyme engineering. Through pathway similarity analysis we classified the pathways and identified the enzymes and precursors that were indispensable for the production of the target molecules. These retrobiosynthesis studies demonstrate the potential of BNICE.ch for discovery, systematic evaluation, and analysis of novel pathways in synthetic biology and metabolic engineering studies.

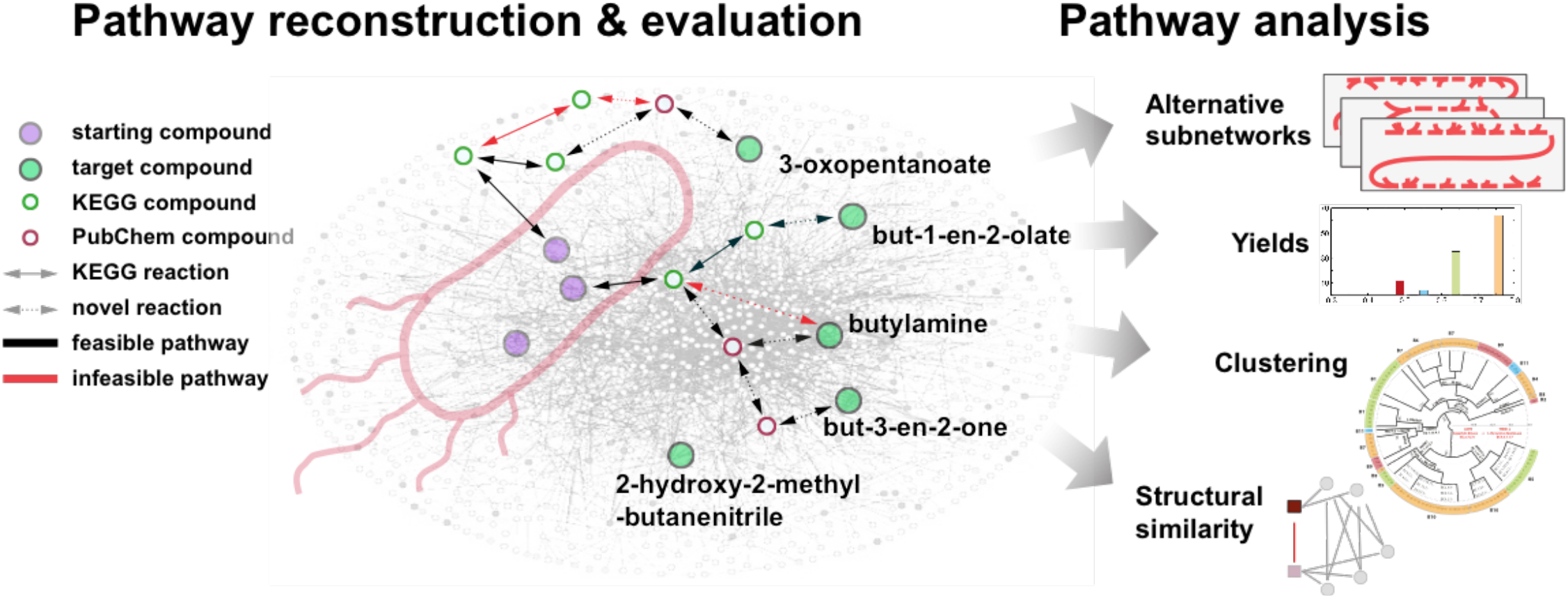
Graphical abstract.

---

Limited reserves of oil and natural gas and the environmental issues associated with their exploitation in the production of chemicals sparked off current developments of processes that can produce the same chemicals from renewable feedstocks using microorganisms.*^1–3^* A fair amount of these efforts focuses on a sustainable production of the 2^nd^ generation biofuels.

Compared to the currently used fossil fuels and bioethanol, these 2^nd^ generation biofuels should provide lower carbon emissions, higher energy density, and should be less corrosive to engines and distribution infrastructures. Recently, a large number of potential candidates for the 2^nd^ generation biofuels has been proposed such as n-butanol*^4^*, isobutanol*^4^*, 2-methyl-1-butanol*^4^*, 3-methyl-1-butanol*^4^*, C13 to C17 mixtures of alkanes and alkenes*^5^*, fatty esters, fatty alcohols*^1^*, and Methyl Ethyl Ketone (MEK)*^6^*.

While some of these compounds were detected in living cells, none was produced by native organisms in appreciable quantities.*^7^* For chemicals whose natural microbial producers are not known, the feasibility of their bioproduction has to be assessed and potential novel biosynthetic pathways for production of these chemicals are yet to be discovered.*^8, 9^* Even when production pathways for target chemicals are known, it is important to find alternatives in order to further reduce cost and greenhouse gas emissions, and as well to avoid possible patent issues.

Computational approaches provide valuable assistance in the design of novel biosynthetic pathways because they allow exhaustive generation of alternative novel biosynthetic pathways and evaluation of their properties and prospects for producing target chemicals.*^9^* For instance, computational tools can be used to assess, prior to experimental pathway implementation, the performance of a biosynthetic pathway operating in one organism across other host organisms.

There are different computational tools for pathway prediction available in the literature.*^8, 10–19^* An important class of these tools is based on the concept of generalized enzyme reaction rules, which were introduced by Hatzimanikatis and co-workers.*^20, 21^* These rules emulate the functions of enzymes, and they can be used to predict *in silico* biotransformations over a wide range of substrates.*^9^* Most of the implementations of this concept appear in the context of retrobiosynthesis, where the algorithm generates all possible pathways by starting from a target compound and moving backward towards desired precursors.*^3,8–10,14,16,19–25^*

In this study, we used the retrobiosynthesis framework of BNICE.ch*^9, 10,20–25^* to explore the biotransformation space around Methyl Ethyl Ketone (MEK). Besides acetone, MEK is the most commercially produced ketone with broad applications as a solvent for paints and adhesives and as a plastic welding agent.*^26^* MEK shows superior characteristics compared to conventional gasoline and ethanol in terms of its thermo-physical properties, increased combustion stability at low engine load, and cold boundary conditions, while decreasing particle emissions.*^27^* There is no known native microbial producer of MEK, but in the recent studies this molecule was produced in *E. coli^28, 29^* and *S. cerevisiae^6^* by introducing novel biosynthetic pathways. To convert 2,3-butanediol to MEK, Yoneda *et al.^30^* introduced into *E. coli* a B-12 dependent glycerol dehydratase from *Klebsiella pneumoniae.* Srirangan *et al.^29^* expressed in *E. coli* a set of promiscuous ketothiolases from *Cupriavidus necator* to form 3-ketovaleryl-CoA, and they further converted this molecule to MEK by expressing acetoacetyl-CoA:acetate/butyrate:CoA transferase and acetoacetate decarboxylase from *Clostridium acetobutylicum.* In *S. cerevisiae*, Ghiaci *et al.^6^* expressed a B12-dependent diol dehydratase from *Lactobacillus reuteri* to convert 2,3-butanediol to MEK. Alternatively, hybrid biochemical/chemical approaches were proposed where precursors of MEK were biologically produced through fermentations and then catalytic processes were used to produce MEK.*^30, 31^*, We used the BNICE.ch algorithm to generate a network of potential biochemical reactions around MEK, and we identified 159 biochemical and 1’ 166 chemical compounds one reaction step away from MEK (Table S1 – Supporting Information). We considered as biochemical compounds the ones that we found in the KEGG^32, 33^ database, and as chemical compounds the ones that we found in the PubChem^34, 35^ but not in the KEGG database. A set of 154 compounds appeared in both databases. Out of these 1’325 compounds, 2-hydroxy-2-methyl-butanenitrile (MEKCNH) was the only KEGG compound connected to MEK through a KEGG reaction (KEGG R09358). For further study, we chose MEKCNH along with three KEGG compounds: 3-oxopentanoate (3OXPNT), but-3-en-2-one (MVK) and butylamine (BuNH_2_), and one PubChem compound: 1-en-2-olate (1B2OT). The latter four compounds were chosen based on two important properties: (i) their simple chemical conversion to MEK, e.g., 3OXPNT spontaneously decarboxylates to MEK; and (ii) their potential use as precursor metabolites to further produce a range of other valuable chemicals.*^36–38^* MVK can be converted to MEK by a 2-enoate reductase from *Pseudomonas putida, Kluyveromyces lactis* or *Yersinia bercovieri,^39^* however, these reactions are not catalogued in KEGG. Similarly, 3OXPNT can be decarboxylated to MEK by acetoacetate decarboxylase from *Clostridium acetobutylicum*.*^29^* In contrast, there are no known enzymes that can convert 1B2OT and BuNH_2_ to MEK.

We have reconstructed all possible novel biosynthetic pathways (3’679’610 in total) up to a length of 4 reaction steps from the central carbon metabolites of *E. coli* towards the 5 compounds mentioned above. We evaluated the feasibility of these 3’679’610 pathways with respect to the mass and energy balance, and we found 18’622 thermodynamically feasible pathways which we further ranked with respect to their carbon yields. We identified the metabolic subnetworks that were carrying fluxes when the optimal yields were attained, and we determined the minimal sets of precursors and the common routes and enzymes for production of the target compounds.

## Results and Discussion

### Generated metabolic network around Methyl Ethyl Ketone

We used the retrobiosynthesis algorithm of BNICE.ch to reconstruct the biochemical network around MEK. BNICE.ch*^9, 10,20–25^* is a computational framework that takes advantage of the biochemical knowledge derived from the thousands of known enzymatic reactions to predict all possible biotransformation pathways from known compounds to desired target molecules. We applied BNICE.ch and generated all compounds and reactions that were up to five reaction steps away from MEK (Figures 1a and 1b).

**Figure 1.**
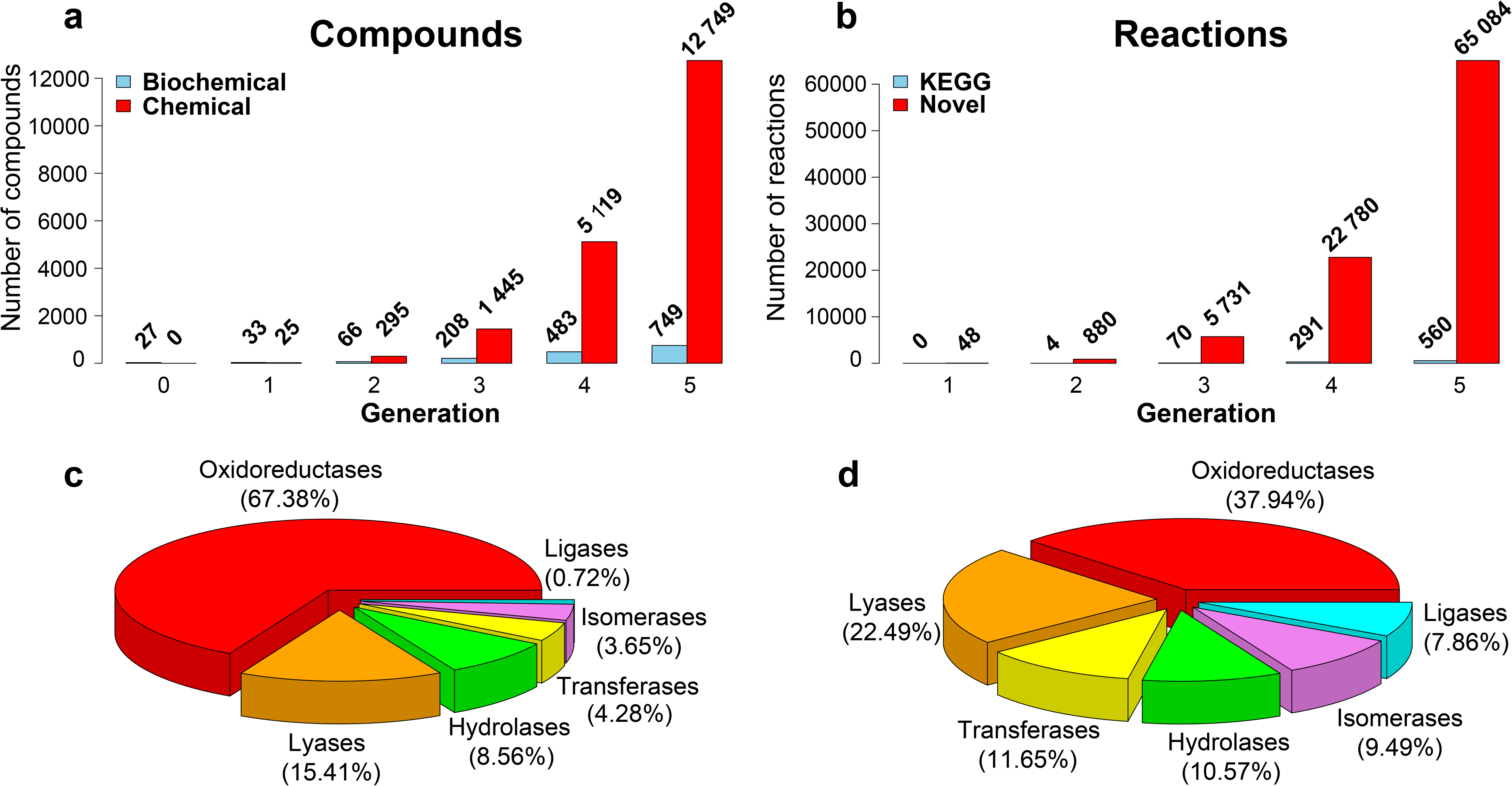
The growth of the generated metabolic network over 5 generations. **(a)** The BNICE.ch retrobiosynthesis algorithm generated 749 biochemical (blue) and 12’749 chemical (red) compounds. **(b)** Generated compounds participated in 560 KEGG (blue) and 65’084 novel (red) reactions. Categorization of the predicted reactions **(c)** and utilized generalized enzyme reaction rules **(d)** on the basis of their Enzymatic Commission^40^, EC, classification.

To start the reconstruction procedure, we provided the initial set of compounds that contained 26 cofactors along with MEK (Table S2 – Supporting Information). In the first BNICE.ch generation, we produced 6 biochemical and 25 chemical compounds connected through 48 reactions to MEK. Interestingly, among these reactions were also the ones proposed by Yoneda *et al.^30^*, Srirangan *et al.^29^* and Ghiaci *et al.^6^* After five generations, a total of 13’498 compounds were generated (Figure 1a). Out of these, 749 were biochemical and the remaining 12’749 were chemical compounds. We could also find 665 out of the 749 biochemical compounds in the PubChem database. All generated compounds were involved in 65’644 reactions, out of which 560 existed in the KEGG database and the remaining 65’084 were novel reactions (Figure 1b). A large majority of the predicted reactions (67%) were oxidoreductases, 15.4% were lyases, 8.6% were hydrolases, 4.3% transferases, 3.6% isomerases and only 0.72% ligases (Figure 1c). Out of 361 bidirectional generalized enzyme reaction rules of BNICE.ch, 369 were required to generate the metabolic network around MEK with the size of 5 reaction steps. As expected from the statistics on the predicted reactions, most of these rules (38%) described the oxidoreductase biotransformation (Figure 1d).

Though MEK participated in a total of 1’551 reactions (Supporting information – Table S3) only one reaction, which connected MEK to MEKCNH, was catalogued in the KEGG database (KEGG R09358). These 1’551 reactions connected MEK to 1’325 compounds (159 biochemical and 1’ 166 chemical), which could be potentially used as MEK precursors (Table S1 – Supporting Information). Reaction steps for a biochemical production of MEK from the five precursors (3OXPNT, MVK, BuNH_2_ 1B2OT, and MEKCNH) together with their most similar KEGG reactions can be consulted in Table S4 – Supporting information.

### Pathway reconstruction towards five target compounds

In the pathway reconstruction process, we used as starting compounds 157 metabolites selected from the generated network, which were identified as native *E. coli* metabolites using the *E. coli* genome-scale model iJO1366^41^ (Table S5 – Supporting Information). We performed an exhaustive pathway search on the generated metabolic network, and we reconstructed 3’679’610 pathways towards these five target compounds with pathway lengths ranging from 1 up to 4 reaction steps (Table 1). The reconstructed pathways combined consist of 37’448 reactions, i. e., 57% of the 65’644 reactions reproduced from the BNICE.ch generated metabolic network.

**Table 1.**
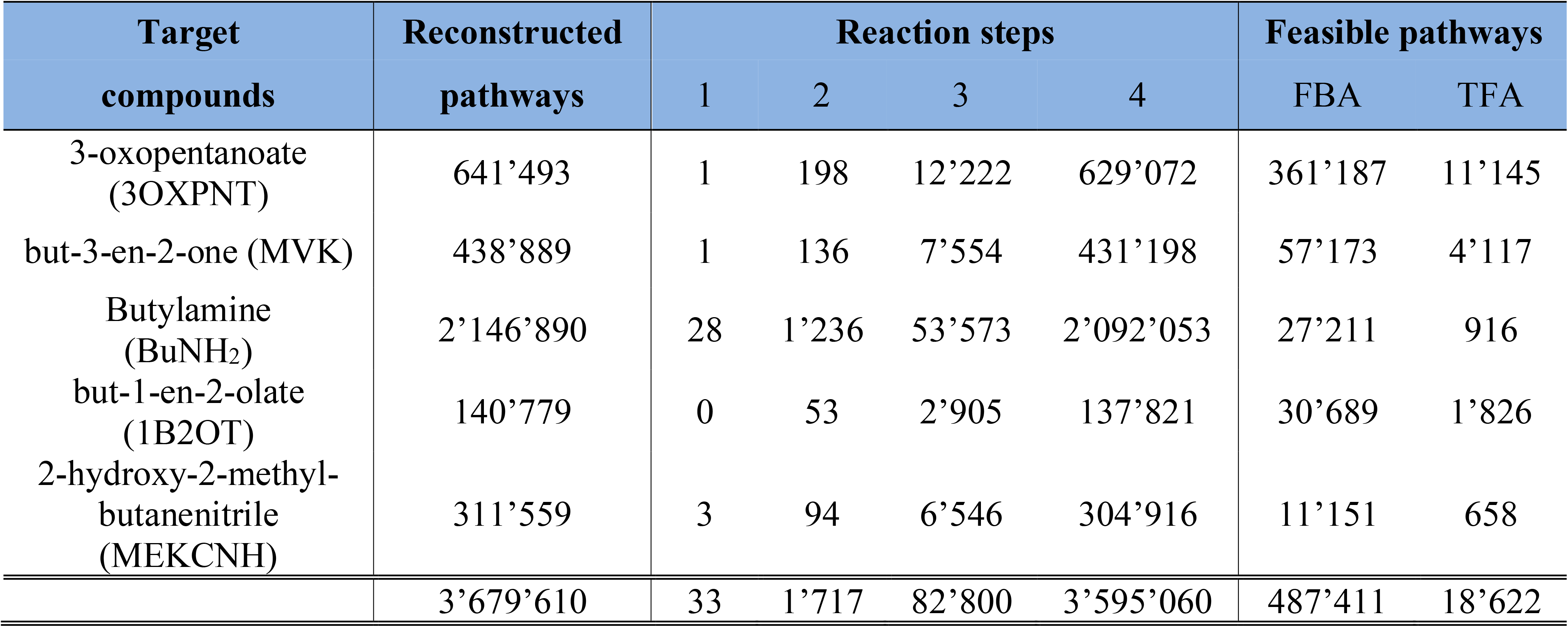
Reconstructed pathways towards five target compounds.

| Target compounds | Reconstructed pathways | Reaction steps |  |  |  | Feasible pathways |  |
| --- | --- | --- | --- | --- | --- | --- | --- |
|  |  | 1 | 2 | 3 | 4 | FBA | TFA |
| 3-oxopentanoate (3OXPNT) | 641'493 | 1 | 198 | 12'222 | 629'072 | 361'187 | 11'145 |
| but-3-en-2-one (MVK) | 438'889 | 1 | 136 | 7'554 | 431'198 | 57'173 | 4'117 |
| Butylamine (BuNH<sub>2</sub>) | 2'146'890 | 28 | 1'236 | 53'573 | 2'092'053 | 27'211 | 916 |
| but-1-en-2-olate (1B2OT) | 140'779 | 0 | 53 | 2'905 | 137'821 | 30'689 | 1'826 |
| 2-hydroxy-2-methyl-butanenitrile (MEKCNH) | 311'559 | 3 | 94 | 6'546 | 304'916 | 11'151 | 658 |
|  | 3'679'610 | 33 | 1'717 | 82'800 | 3'595'060 | 487'411 | 18'622 |

More than 58% of the discovered pathways were towards BuNH_2_ while only 3.8% of the reconstructed pathways were towards 1B2OT, which was the only PubChem target compound (Table 1). Only 33 reconstructed pathways were of length one, and 28 out of them were towards BuNH2 and none towards 1B2OT. The majority of reconstructed pathways (> 97%) were of length four. These results suggest that the biochemistry of enzymatic reactions favors smaller changes of a molecule structure over several steps.

### Evaluation of reconstructed pathways

We performed a series of studies of the 3’679’610 generated pathways to assess their biological feasibility and performance (Methods). The feasibility of the pathways depends on the metabolic network of the chassis organism. Therefore, we embedded each of the reconstructed pathways in the *E. coli* genome-scale model iJO1366 and performed flux balance analysis (FBA)*^42^* and thermodynamics-based flux analysis (TFA)*^43–47^*. The directionality of the reactions is an important factor in FBA and TFA*^45^*, and in our studies, unless stated otherwise, for FBA and TFA we applied the *C1* constraints on reaction directionalities where we constrained the reactions that involve CO2 to operate in the decarboxylation direction (Methods).

#### Flux balance analysis

We used FBA as a prescreening method to reject pathways that were incompatible with the host organism (Methods). If an FBA model formed by embedding a pathway in iJO1366 can produce the target compound, then the pathway is considered as FBA feasible. Out of all reconstructed pathways, only 13.24% (487’411) were FBA feasible (Table 1). Though the largest number of reconstructed pathways were towards BuNH_2_, only 1.27% (27’211) of these were FBA feasible. The number of FBA feasible pathways for MEKCNH was also low (3.59%). In contrast, more than 56% of pathways towards 3OXPNT were FBA feasible.

#### Thermodynamics-based flux analysis

We used TFA to identify 18’622 thermodynamically feasible pathways (0.5% of all generated pathways, or 3.8% of the FBA feasible pathways). A pathway is considered TFA feasible if a TFA model formed by embedding the pathway in iJO1366 can produce the target compound under thermodynamic constraints. The set of TFA feasible pathways involved 3’ 166 unique reactions. These results demonstrate that TFA is important for pathway evaluation and screening.

We found BuNH_2_ to have the lowest rate of TFA feasible pathways with 0.04% of reconstructed pathways being TFA feasible (Table 1). The highest rate of TFA feasible pathways was again for 3OXPNT (1.74 %). The shortest TFA feasible pathways consisted of 2 reaction steps (21 pathways), whereas a majority of TFA feasible pathways had length 4 (Table 2). All pathways contained novel reaction steps, and only 19 pathways had one novel reaction step (Table 2). All of these 19 pathways were towards MVK, and they all had as intermediates 2-acetolactate and acetoin. The final reaction step converting acetoin to MVK was novel in all of them.

**Table 2.**
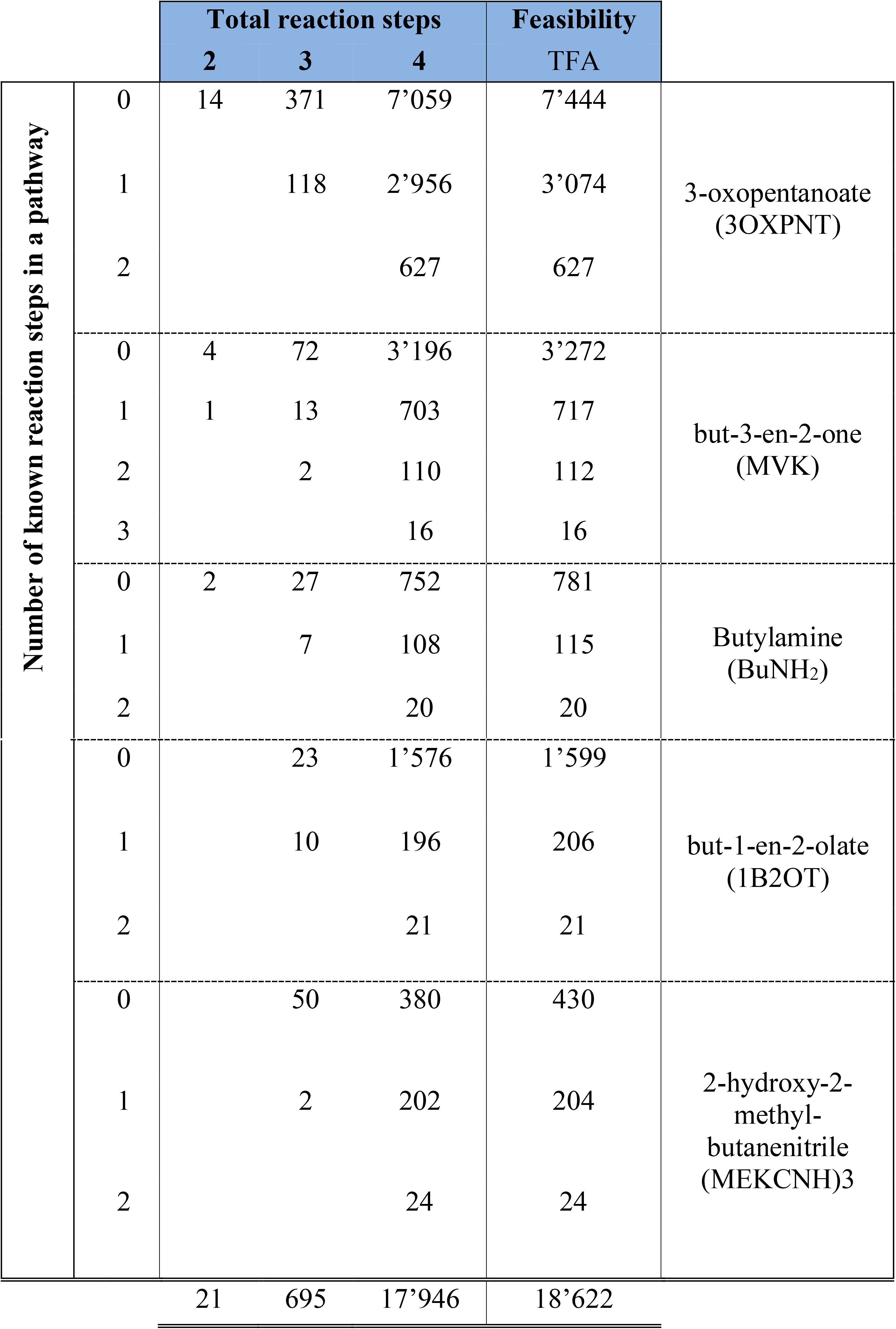
Number of known reaction steps versus all reaction steps in the predicted pathways. Table should be read as follows: e.g., among the predicted pathways of length 3 towards 3OXPNT, there were 371 pathways with all novel reaction steps (no known steps), 118 pathways with 1 known and 2 novel reaction steps, and no pathways with 2 known and 1 novel reaction steps. Pathways with one novel reaction step are marked in red. All shown pathways are TFA feasible.

#### Yield analysis

We used TFA to assess the production yield of the feasible pathways from glucose to the target compounds (Table S6 – Supporting Information). We identified pathways for all target compounds that could operate without a loss of carbon from glucose. More than a half of the pathways towards 3OXPNT (57%) could operate with the maximum theoretical yield of 0.774 g/g, i.e., 1Cmol/1Cmol (Figure 2). In contrast, only 4% (25 out of 658) pathways towards MEKCNH could operate with the maximal theoretical yield of 0.66 g/g (Table S6 – Supporting Information). We found that pathway yields were distributed into several distinct sets rather than being more spread and continuous, i.e., we obtained eleven sets for 3OXPNT, four sets for MEKCNH, 11 sets for BuNH_2_ nine sets for 1B2OT and ten sets for MVK (Table S6 – Supporting Information). Interestingly, a discrete pattern in pathway yields was also observed in a similar retrobiosynthesis study for the production of mono-ethylene glycol in *Moorella thermoacetica* and *Clostridium ljungdahlii.^48^*

**Figure 2.**
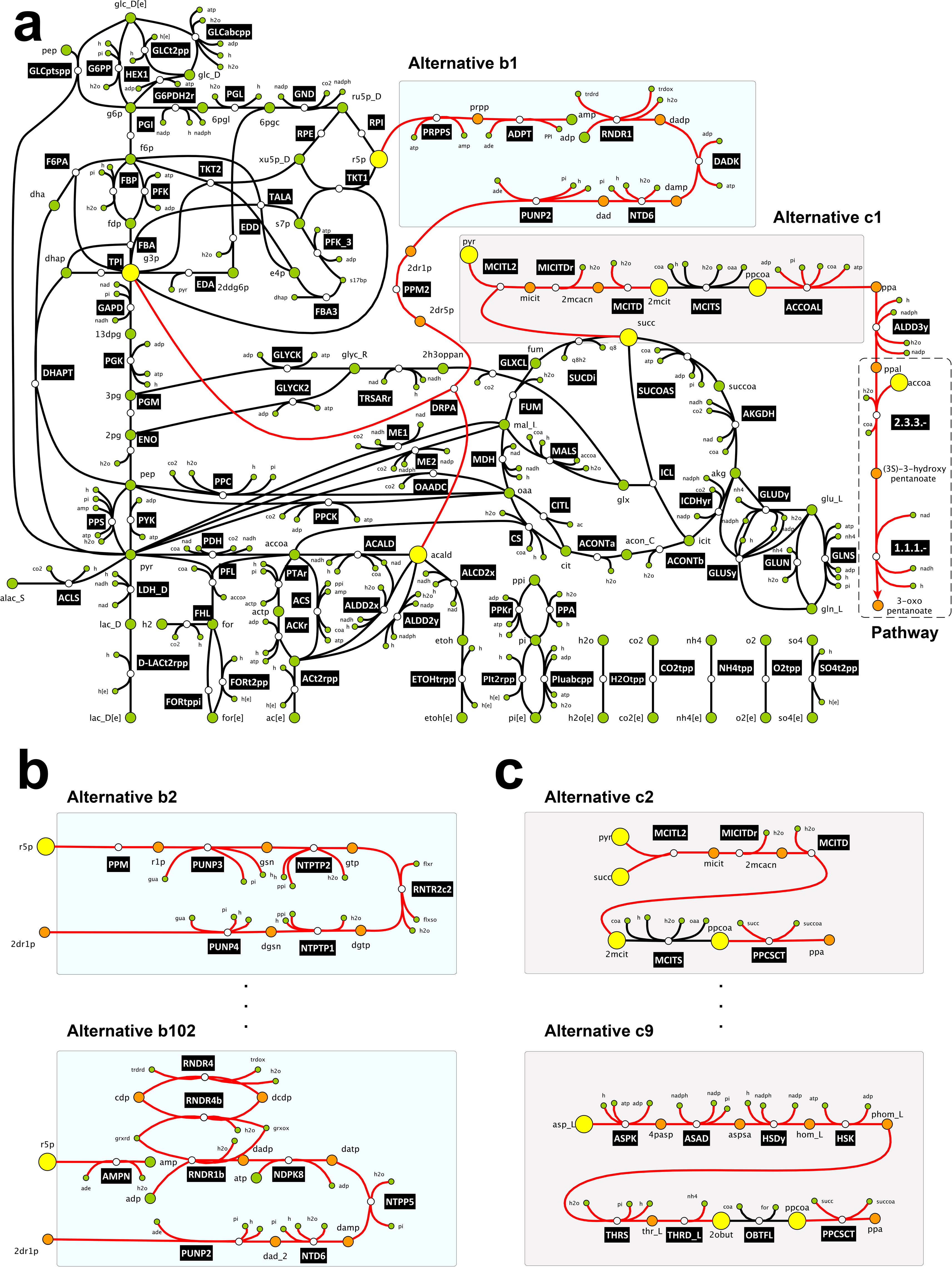
Alternative ways of producing 3OXPNT from glucose. (a) Schematic representation of the metabolic network producing 3OXPNT from glucose. Reactions pertaining to the core metabolic (black) and the active anabolic (red) subnetworks. Metabolites of the core metabolic (green) and the active anabolic (orange) subnetworks together with core precursors (yellow), i.e., metabolites that connect the core and active anabolic subnetworks. (b) Alternative pathways connecting ribose-5-phosphate, r5p, with 2-deoxy-D-ribose-1-phosphate, 2dr1p. (c) Alternative pathways connecting the core metabolites with propanal, Ppal.

#### Analysis of alternative assumptions on reaction directionalities

Since we found that the directionality of reactions in the network impacts yields, we investigated how the type of alternative constraints *C2* affected the yield distribution (Methods). The *C2* constraints contain the preassigned reaction directionalities from the iJO1366 model together with the *C1* constraints. As expected, these additional constraints reduced flexibility of the metabolic network and some pathways even became infeasible (Table S7 – Supporting Information). With the *C2* constraints, the yields were in general reduced and their distribution was more spread compared to the one obtained using the *C1* constraints. For example, we found with both sets of constraints three alternative pathways for the production of 3OXPNT from acetate via two intermediate compounds: 2-ethylmalate and (3S)-3-hydroxypentanoate. The three alternative pathways had three different cofactor pairs in the final reaction step that converts (3S)-3-hydroxypentanoate to 3OXPNT (Figure F1 – Supporting Information). With the *C1* constraints, the three pathways had an identical yield of 0. 642 g/g. In contrast, with the *C2* constraints, the pathway with NADH/NAD cofactor pair in the final step had a yield of 0.537 g/g, the one with NADPH/NADP had a yield of 0.542 g/g, and the one with H2O2/H20 had a yield of 0.495 g/g. These differences in yields are a consequence of the different costs of cofactor production upon adding supplementary constraints.

These results highlight the importance of the choice of constraints in FBA and TFA as they can influence our conclusions on reaction directionalities. Besides, the reaction directionalities have a critical impact on network properties such as gene essentiality or yields.*^45^* This suggests that particular caution should be exercised when using “off-the-shelf’ models as some of them have *ad hoc* pre-assigned directionalities.*^45–47^* Additionally, this indicates that there is a need for revisiting assumptions on reaction directionalities in the current genome-scale reconstructions. This task can be performed by integrating thermodynamic constraints in metabolic networks and thus allowing for systematical assigning of reaction directionalities.*^45^* However, for an accurate estimation of the reaction directionalities using thermodynamics, it is crucial to consider the contribution of metabolite activities to the Gibbs free energy of reactions instead of using only the standard values.*^46,47^* Since metabolite activities are proportional to metabolite concentrations,*^49^* this further emphasizes the importance of integrating metabolomics data.

#### BridgIT analysis

For each novel reaction from the feasible pathways, we identified the most similar KEGG reaction whose gene and protein sequences were assigned to the novel reaction (Methods). The BridgIT^50^ results can be consulted at http://lcsb-databases.epfl.ch/GraphList/ProjectList upon subscription.

### Identification and analysis of anabolic subnetworks capable of synthesizing target molecules

In pathway reconstruction, we identified the sequence of the main reactions required to produce the target molecules from precursor metabolites in the core network. However, these reactions require additional co-substrates and cofactors that should become available from the rest of the metabolism. In addition, these reactions produce also side products and cofactors that must be recycled by the genome-scale metabolic network in order to have a biologically feasible and balanced subnetwork for the production of the target molecules. Therefore, we identified the active metabolic subnetworks required to synthesize the corresponding target molecule (Methods). The active metabolic subnetworks were then divided into the *core metabolic network*, which included central carbon metabolism pathways*^51, 52^*, and the active *anabolic subnetwork* (Figure 2a, and Methods). On average there were more than three alternative anabolic subnetworks per pathway due to the redundant topology of metabolism (Table 3). For example, we identified 35’013 alternative anabolic subnetworks for 11’ 145 feasible pathways towards 3OXPNT. Overall, for the 18’622 TFA feasible pathways, 55’788 active anabolic subnetworks were identified.

**Table 3.** Alternative anabolic subnetworks for 5 target compounds together with their lumped reactions and precursors.

| Target compounds | Feasible pathways | Alternative anabolic subnetworks | Unique lumped reactions | Overlapping sets of precursors | Unique precursors |
| --- | --- | --- | --- | --- | --- |
| 3-oxopentanoate (3OXPNT) | 11'145 | 35'013 | 4'517 | 281 | 40 |
| but-3-en-2-one (MVK) | 4'117 | 10'162 | 1'762 | 126 | 32 |
| Butylamine (BuNH<sub>2</sub>) | 916 | 2'858 | 974 | 102 | 30 |
| but-1-en-2-olate (1B2OT) | 1'826 | 5'339 | 1'536 | 97 | 30 |
| 2-hydroxy-2-methyl-butanenitrile (MEKCNH) | 658 | 2'416 | 792 | 37 | 17 |
|  | 18'622 | 55'788 | 9'581 |  |  |

Next, we computed a lumped reaction for each of the alternative subnetworks (Methods). Similar to previous findings from the analysis of the biomass building blocks in *E. coli,^53^* only 9’581 out of the 55’788 computed lumped reactions were unique (Table 3). This result suggests that the overall chemistry and the cost to produce the corresponding target molecule are the same for many different pathways. Since the cost of producing a target molecule depends of the host organism, this implies that the choice of the host organism is important. On the other hand, the multiple alternative options could also provide useful degrees of freedom for synthetic biology and metabolic engineering design.

The largest diversity in alternative subnetworks per lumped reaction was found for 3OXPNT, where on average more than seven alternative subnetworks had the same lumped reaction (Table 3). In contrast, we observed the smallest diversity for BuNH2 with approximately three alternative subnetworks per lumped reaction (Table 3). An illustrative example of multiple pathways with the same lumped reaction is provided in Figure F2 in the Supporting Information.

Interestingly, the 35’013 active anabolic networks towards the production of 3OXPNT were composed of only 394 unique reactions. Out of these 394 reactions, 132 were common with the pathways leading towards the production of all biomass building blocks (Table S8 – Supporting information) except chorismate, phenylalanine, and tyrosine. This finding suggests that biomass building blocks could be competing for resources with 3OXPNT and that they could affect the production of this compound.

#### Origins of diversity of alternative anabolic subnetworks

To better understand the diversity in alternative anabolic subnetworks, we performed an in-depth analysis of the two-step pathway from acetyl-CoA and propanal to 3OXPNT, which presented the largest number of alternative anabolic networks (185) among all reconstructed pathways (Figure 2a). The smallest anabolic subnetwork of the 185 alternatives consisted of 14 enzymes, whereas the largest one comprised 22 enzymes (Table S9 – Supporting Information). All 185 subnetworks shared five common enzymes: the two enzymes from the reconstructed pathway converting propanal via (3S)-3-hydroxypentanoate to 3OXPNT (with the BNICE.ch assigned third level Enzymatic Commission*^40^*, EC, numbers 2.3.3.- and 1.1.1.-), two enzymes involved in acetyl-CoA production (phosphopentomutase deoxyribose (PPM2), and deoxyribose-phosphate aldolase (DRPA)), and aldehyde dehydrogenase (ALDD3y) that converts propionate to propanal (Figure 2).

The multiplicity of ways to produce acetyl-CoA and propionate contributed to a large number of alternative subnetworks: there were 102 alternative ways of producing acetyl-CoA from ribose-5-phosphate (r5p) via 2-deoxy-D-ribose-1-phosphate (2dr1p) (Figure 2b) and 9 different ways of producing propionate (Figure 2c).

There were two major routes to produce 2dr1p within the 102 alternatives. In the first route with 50 alternatives, r5p is converted either to ribose-1-phosphate (in 31 alternatives) or to D-ribose (in 19 alternatives), which are intermediates in producing nucleosides such as adenosine, guanosine, inosine and uridine. These nucleosides are further converted to deoxyadenosine (dad), deoxyguanosine (dgsn) and deoxyuridine (duri) that are ultimately phosphorylated to 2dr1p. In 26 of the remaining 52 alternatives of the second route, r5p is converted to phosphoribosyl pyrophosphate (prpp), which is followed by a transfer of its phospho-ribose group to nucleotides such as AMP, GMP, IMP and UMP. These nucleotides are then converted to 2dr1p by downstream reaction steps. In the remaining alternatives for the second route, r5p is first converted to AMP in one reaction step, and then to 2dr1p via dad and dgsn.

There were 9 alternative routes to produce propionate. In 4 of these, this compound was produced from pyruvate and succinate (Figure 2a and 2c), in 3 routes it was produced from aspartate (Figure 2c), and in 2 routes it was produced from 3-phosphoglycerate and glutamate.

#### Core precursors of five target compounds

An abundant availability of precursor metabolites is crucial for an efficient production of target molecules.^54^ Here, we defined as *core precursors* the metabolites that connect the core to the active anabolic subnetworks (Figure 2a). We analyzed the different combinations of core precursors that appeared in the alternative subnetworks. Our analysis revealed that the majority of subnetworks were connected to the core network through a limited number of core precursors. We found that all 35’013 alternative subnetworks for the production of 3OXPNT were connected to the core network by 281 sets of different combinations among 40 unique core precursors (Table 3). We ranked these sets based on their number of appearances in the alternative networks. The top ten sets appeared in 24’210 subnetworks, which represented 69% of all identified subnetworks for this compound (Table 4). Moreover, the metabolites from the top set (acetyl-CoA, propionyl-CoA, pyruvate, ribose-5-phosphate, and succinate) were the precursors in 8’510 (24.3%) subnetworks for 3OXPNT (Table 4). Ribose-5-phosphate appeared in 9 out of the top ten sets, and it was a precursor in 32’237 (92%) 3OXPNT producing subnetworks.

**Table 4.**
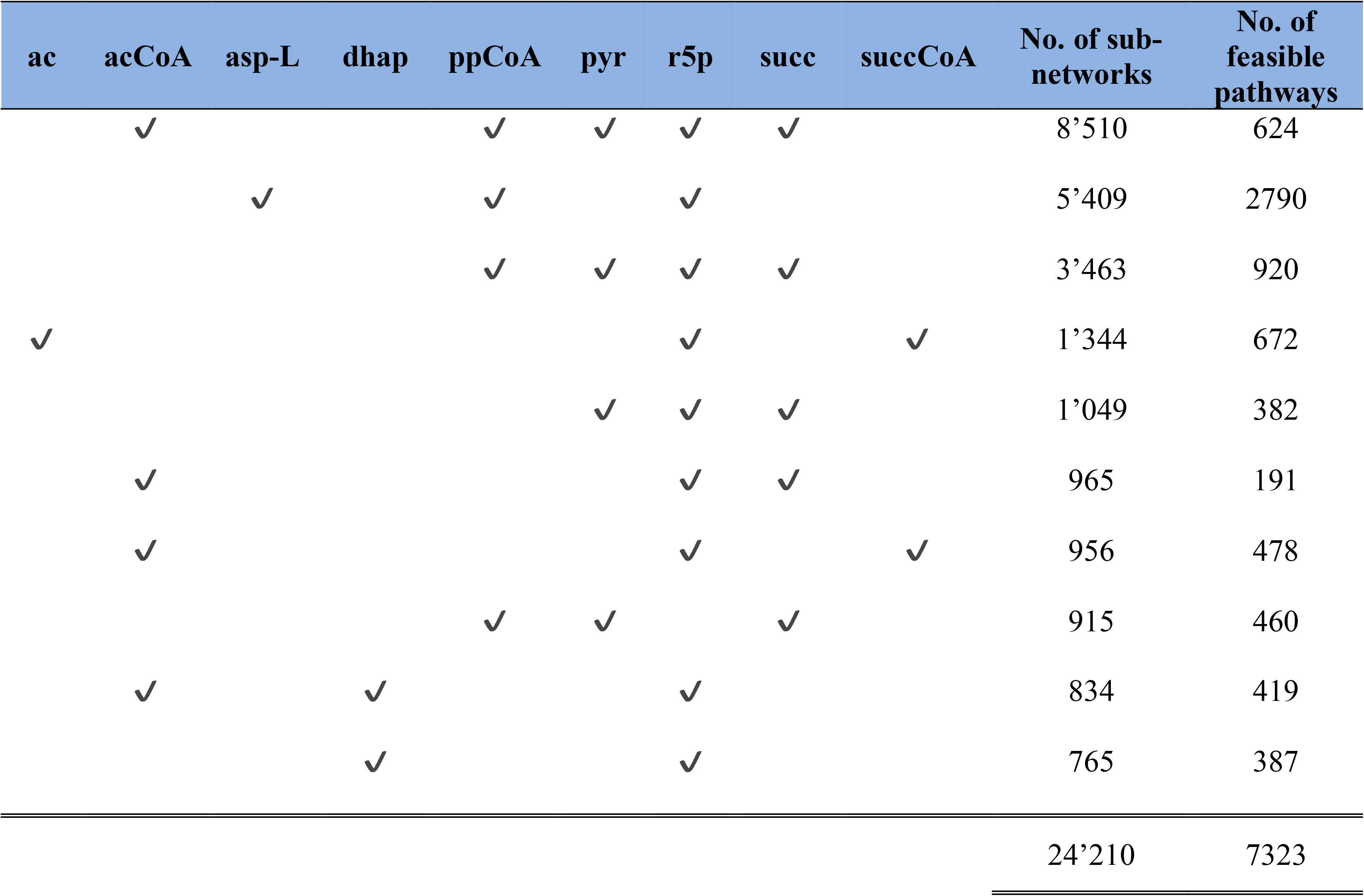
Top ten core precursor combinations for the production of 3OXPNT. Core precursors: acetate (ac), acetyl-CoA (acCoA), aspartate (asp-L), dihydroxyacetone phosphate (dhap), propionyl-CoA (ppCoA), pyruvate (pyr), ribose-5-phosphate (r5p), succinate (succ), succinyl-CoA (succCoA).

### Clustering of feasible pathways

The repeating occurrences of core precursors and lumped reactions in the alternative anabolic subnetworks motivated us to identify common patterns in core precursors, enzymes, and intermediate metabolites required to produce the target molecules. To this end, we used the feasible pathways from acetate to 3OXPNT as the test study, and we performed two types of clustering on these 115 pathways (File M1 – Supporting information).

#### Clustering based on core precursors and byproducts of lumped reactions

We identified 242 alternative anabolic networks for 115 pathways from the test study, and we computed the corresponding 242 lumped reactions (File M1 – Supporting information). We chose the first lumped reaction returned by the solver for each of the 115 pathways, and we clustered the pathways based on the structural similarity between the core precursors and byproducts of the lumped reactions (Methods).

The clustering separated the 115 pathways in eleven groups, B1-B11 (Figure 3a and 3b, Supporting information – Table S10). The main clustering condition among the 115 pathways was the presence or absence of thioesters, such as AcCoA, in the set of core precursors. There were 56 pathways with CoA-related precursors (groups B1-B5) and 59 pathways that did not require CoA (groups B6-B11). The pathways from groups B1-B5 were further clustered subject to the presence of: the precursor succCoA and the byproduct CO2 (group B1); the precursor succCoA and the byproduct malonate (group B2); the precursor ppCoA (group B3); the precursors acCoA and ppCoA (group B4); and the precursor acCoA (group B5). The pathways that did not require CoA were further clustered depending on if they had as precursors dhap (groups B6 and B7) or formate (B8) or not (B9-B11). The clustering results for the complete set of 242 lumped reactions are provided in the Supporting information (Figure F3).

**Figure 3.**
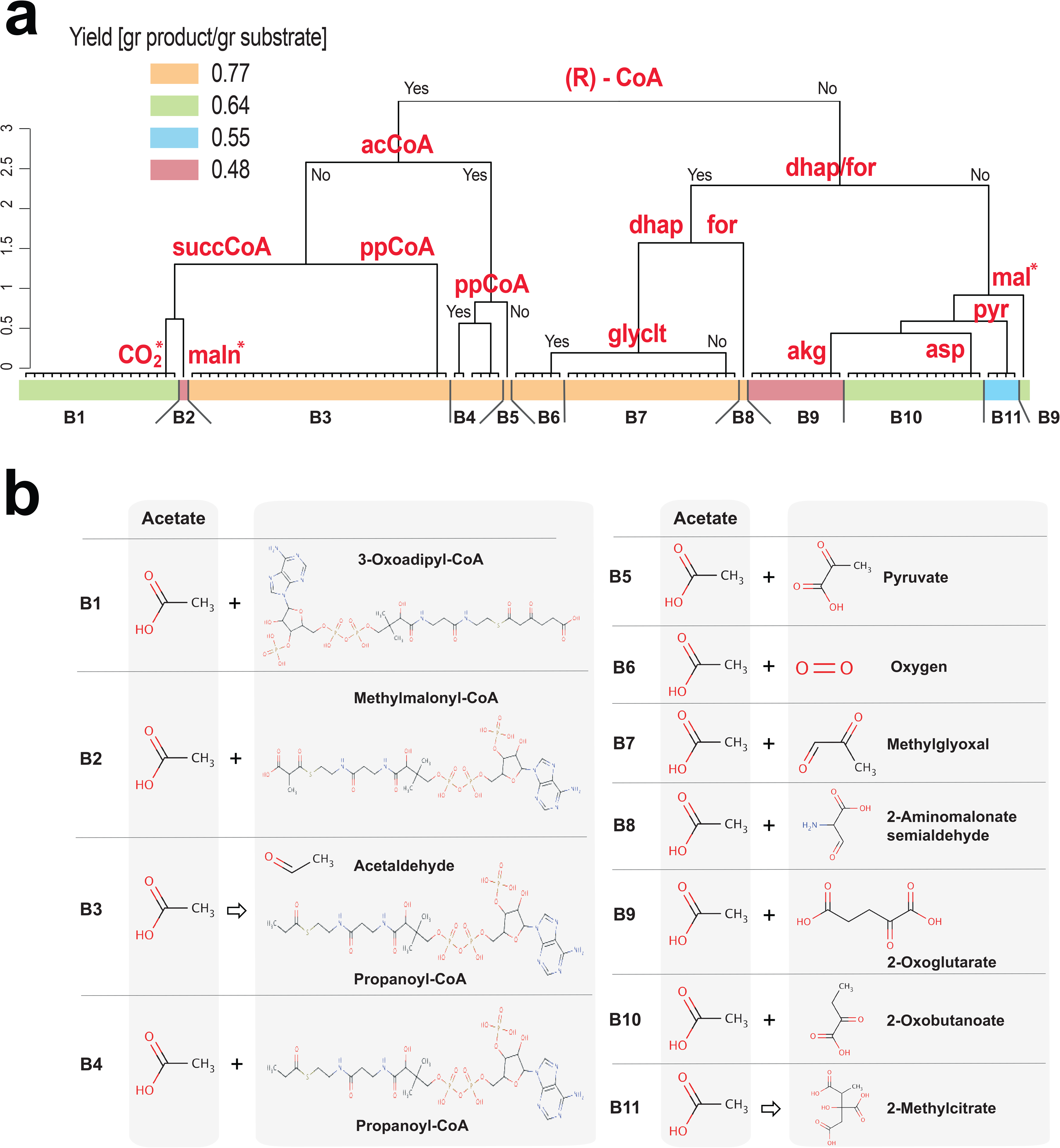
Clustering of the 115 reconstructed pathways from acetate to 3OXPNT. (a) Pathways were classified in eleven groups (B1-B11) based on core precursors and byproducts of their lumped reactions. The byproducts are denoted with an asterisk (*). The pathway yields were consistent within each of the groups and distinctly separated between them. The color-coding of the groups corresponds to the yields of the involved pathways. (R)-CoA denotes the group of thioesters. Abbreviations: 2-oxoglutarate (akg), acetyl-CoA (acCoA), aspartate (asp), dihydroxyacetone phosphate (dhap), formate (for), glycolate (glyclt), malate (mal), malonate (maln), propionyl-CoA (ppCoA), pyruvate (pyr), succinyl-CoA (succCoA). (b) Nine out of eleven groups of reconstructed pathways (B1-B2 and B4-B10) are characterized by the core precursors and their co-substrates in the first reaction step of the pathways. Group B3 is characterized by intermediate metabolites acetaldehyde and propionyl-CoA involved in the novel reaction step EC 2.3.1.-. Group B11 pathways have as an intermediate 2-methylcitrate.

In general, we expect the set of precursors and byproducts to affect the pathway yield. Interestingly, the clustering based on core precursors and byproducts of lumped reactions also separated distinctly the pathways based on their yields (Figure 3a). Pathways that have ppCoA, acCoA, dhap, and for as precursors (groups B3-B8) have a maximal theoretical yield of 0.774 g/g. Despite sharing the first reaction step in which acetate reacted with 2-oxoglutarate to create 2-hydroxybutane 1-2-4-tricarboxylate, the pathways from group B9 were split in two groups with different yields (Figure 3a). These two groups differed in the sequences of reactions involved in the reduction of 2-hydroxybutane 1-2-4-tricarboxylate, a 7-carbon compound, to 3OXPNT. In 11 pathways, the yield was 0.483 g/g due to a release of two CO2 molecules, whereas in one pathway the yield was 0.644 g/g due to malate being created as a side-product and recycled back to the system. The pathway from group B2, with succCoA as a precursor and maln as a byproduct, together with the 11 pathways from group B9 had the lowest yield (0.483 g/g) from the set of examined pathways (Figure 3a).

The clustering also provided insight into the different chemistries behind the analyzed pathways. For most of the pathways, i.e., the ones classified in groups B1-B2 and B4-B10, there was a clear link between the core precursors and co-substrates of acetate in the first reaction step of the pathways (Figure 3b). For example, the pathways from the group B1 have a common first reaction step (EC 2.8.3.-) that converts acetate and 3-oxoadipyl-CoA to 3-oxoadipate (Figure 3b). The clustering grouped these pathways together because succCoA was the core precursor of 3-oxoadipyl-CoA through 3-oxoadipyl-CoA thiolase (3-OXCOAT). Moreover, 3-oxoadipate, a 6-carbon compound, was converted in downstream reaction steps to 3OXPNT, a 5-carbon compound, and one molecule of CO2 through 18 alternative routes. Similarly, in the single pathway of group B2 the co-substrate in the first reaction step was (S)-methylmalonyl-CoA, which was produced from succCoA through methylmalonyl-CoA mutase (MMM). This enzyme, also known as sleeping beauty mutase, is a part of the pathway converting succinate to propionate in *E. coli.^55^* Malonate (maln), a 2-carbon compound, was released in the first reaction step, which resulted in a low yield of this pathway (Figures 3a and 3b).

Pathways from group B3 utilized different co-substrates, such as ATP and crotonoyl-CoA, along with acetate to produce acetaldehyde in the first reaction step. All these pathways shared a common novel reaction step with acetaldehyde and propionyl-CoA as substrates (EC 2.3.1.-).

Finally, group B11 contained the pathways with the intermediate 2-methylcitrate, which was produced from pyruvate (pyr).

The presented clustering analysis has been shown to be very powerful in identifying the features of the large number of pathways. The classification can further guide us to identify the biochemistry responsible for the properties of pathways. Such deeper understanding can provide further assistance for the design and analysis of novel synthetic pathways.

#### Clustering based on involved enzymes

Although the clustering based on the core precursors and byproducts provided an insight of the chemistry underlying the production of 3OXPNT from acetate, lumped reactions conceal the identity of the enzymes involved in the active anabolic subnetworks. We analyzed the 115 active subnetworks corresponding to 115 pathways (File M1 – Supporting information), and we found that five enzymes were present in all of them: AMP nucleosidase (AMPN), 5’-nucleotidase (NTD6), purine-nucleoside phosphorylase (PUNP2), PPM2 and DRPA, which participated in the production of acetaldehyde from r5p (Figure 4b).

**Figure 4.**
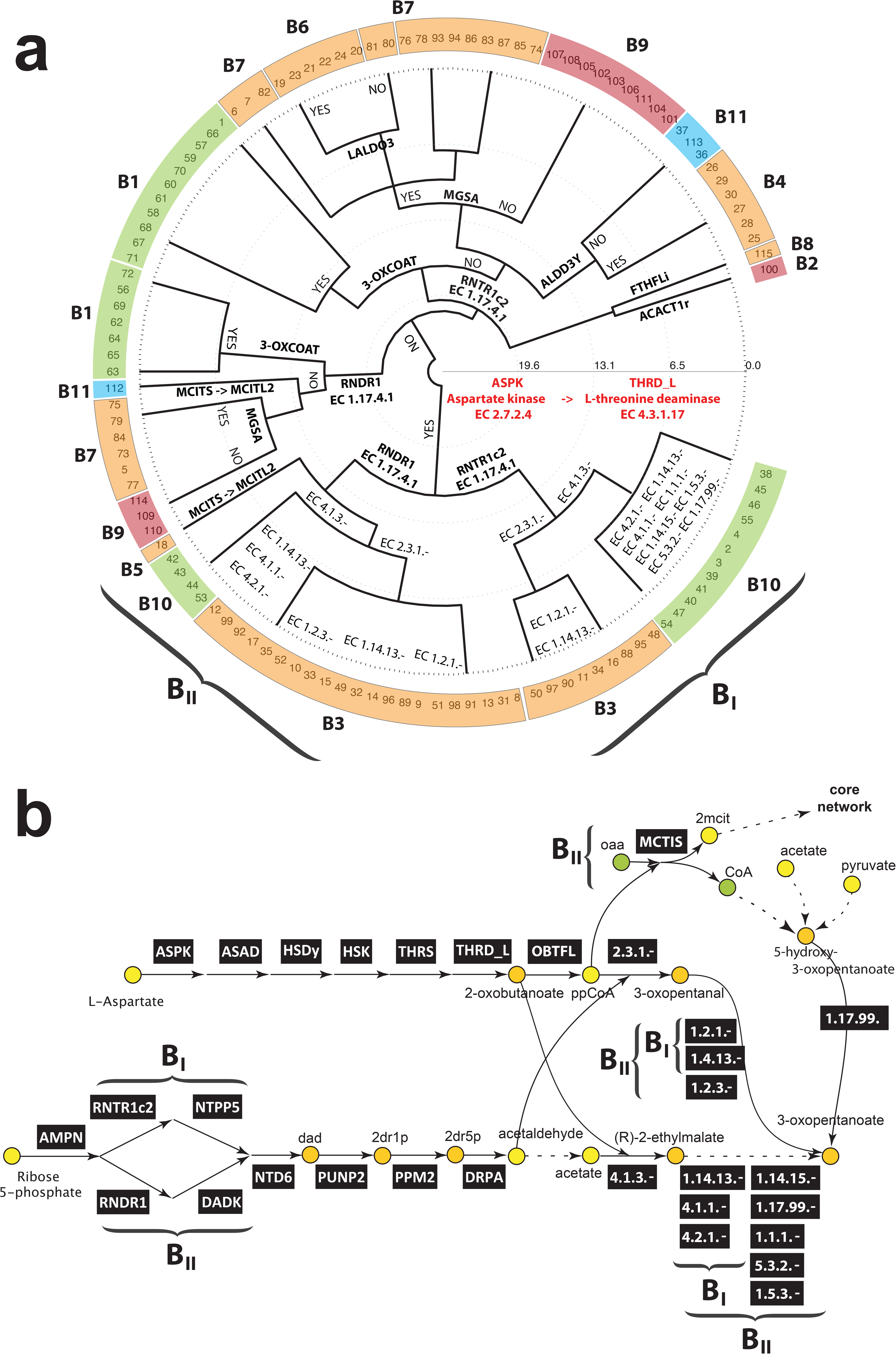
Clustering of 115 active subnetworks corresponding to 115 reconstructed pathways from acetate to 3OXPNT. **(a)** Subnetworks were clustered based on enzymes they involved. The groups B1-B11 are color-coded as in Figure 3a. (**b**) Structure of 47 subnetworks containing a sequence of six enzymes starting with aspartate kinase (ASPK) and ending with L-threonine deaminase (THRD_L) (groups BI and BII in a). The core metabolites are marked in green, the core precursors in yellow, while the metabolites from the active anabolic networks are marked in orange.

To find common enzyme routes in these subnetworks, we performed a clustering based on the structural similarity between their constitutive reactions (Methods). The clustering separated 115 subnetworks in two groups depending on the existence (47 subnetworks) or not (68 subnetworks) of a sequence of six enzymes starting with aspartate kinase (ASPK) and ending with L-threonine deaminase (THRD_L), whose product 2-oxobutanoate was converted downstream to 3OXPNT (Figures 4a and 4b).

Both groups were further clustered based on a set of enzymes required to produce deoxyadenosine and the downstream metabolite acetaldehyde (Figures 4a and 4b). The first subgroup of enzymes, i.e. ribonucleoside-diphosphate reductase (RNDR1), deoxyadenylate kinase (DADK) and NTD6, converted adp to deoxyadenosine. In the second subgroup, atp was transferred to deoxyadenosine via ribonucleoside-triphosphate reductase (RNTR1c2), nucleoside triphosphate pyrophosphorylase (NTPP5) and NTD6 (Figure 4b). Then, for both subgroups, deoxyadenosine was converted to 2-deoxy-D-ribose 5-phosphate (2dr5p) that was further transformed to acetaldehyde via PPM2 and DRPA (Figures 2 and 4b).

The clustering based on enzymes allowed us to identify enzymatic routes corresponding to different yields (Figure 4a, and Supporting information – Table S6). For example, all pathways that include ASPK and novel reaction steps with the third level EC class 2.3.1.- and 1.2.1.-(group B3), would provide the maximal theoretical yield of 0.774 g/g (Figure 4a). Similarly, pathways that contain ALDD3Y (group B4), methylglyoxal synthase (MGSA) (groups B6 and B7), and ASPK, RNDR1, and methylisocitrate lyase (MCITL2) (group B5), would also provide the maximal theoretical yield. In contrast, the clustering also permitted us to identify key enzymes participating in pathways with a reduced yield. For example, pathways that contained 3-OXCOAT had a yield of 0.644 g/g.

Furthermore, the clustering based on enzymes allowed us to clarify the link between the precursors and the corresponding sequence of enzymes that needed to be active for producing the target molecule. For example, pathways from group B1, which had succCoA as a core precursor and CO2 as a byproduct, had the common reaction step 3-OXCOAT (Figure 4a). Similarly, all pathways from group B4 with core precursors ppCoA and acCoA contained ALDD3Y.

### Ranking of biosynthetic pathways and recommendations

We further ranked the corresponding feasible pathways according to number of reaction steps and enzymes that could be directly implemented or needed to be engineered, their yield, and the BridgIT score of the novel reaction steps (Methods). As we saw earlier (e.g. in Supporting information, Table S6), there are several distinct maximum yield values that can be achieved with all these alternatives rather than a continuous distribution of yields. The clustering analysis suggests that the reason for the discreet distribution is the loss of the carbon atoms in specific steps along the pathways. We obtained the top candidate pathways for each of the target molecules that were likely to produce these compounds with economically viable yields (Supporting information – Tables S11-S15). For each of the target molecules, the highest ranked candidate pathways could operate with their maximum theoretical yields (Figure 5). Furthermore, the BridgIT results suggest that the novel reactions appearing in these pathways can be catalyzed by the known enzymes (Figure 5). The highest ranked candidate pathway among all feasible pathways was from pyruvate to 3OXPNT, and it consisted of two novel reactions of the third level EC class 2.3.3- and 1.1.1.- (Figure 5a). The BridgIT^50^ analysis identified KEGG R00472 as the most similar reaction to 2.3.3.-. KEGG reports that R00472 can be catalyzed by EC 2.3.3.9. Similarly, KEGG R01361 was identified as the most similar to 1.1.1-, and according to the KEGG database this reaction is catalyzed by EC 1.1.1.30. Interestingly, there is a reaction that involves CO2 in the top pathways for MVK and MEKCNH (Figure 5b and 5e). Although in the literature this reaction is reported to operate in the decarboxylation direction, TFA allows it to operate in the opposite direction as well. Without TFA this information would stay hidden.

**Figure 5.**
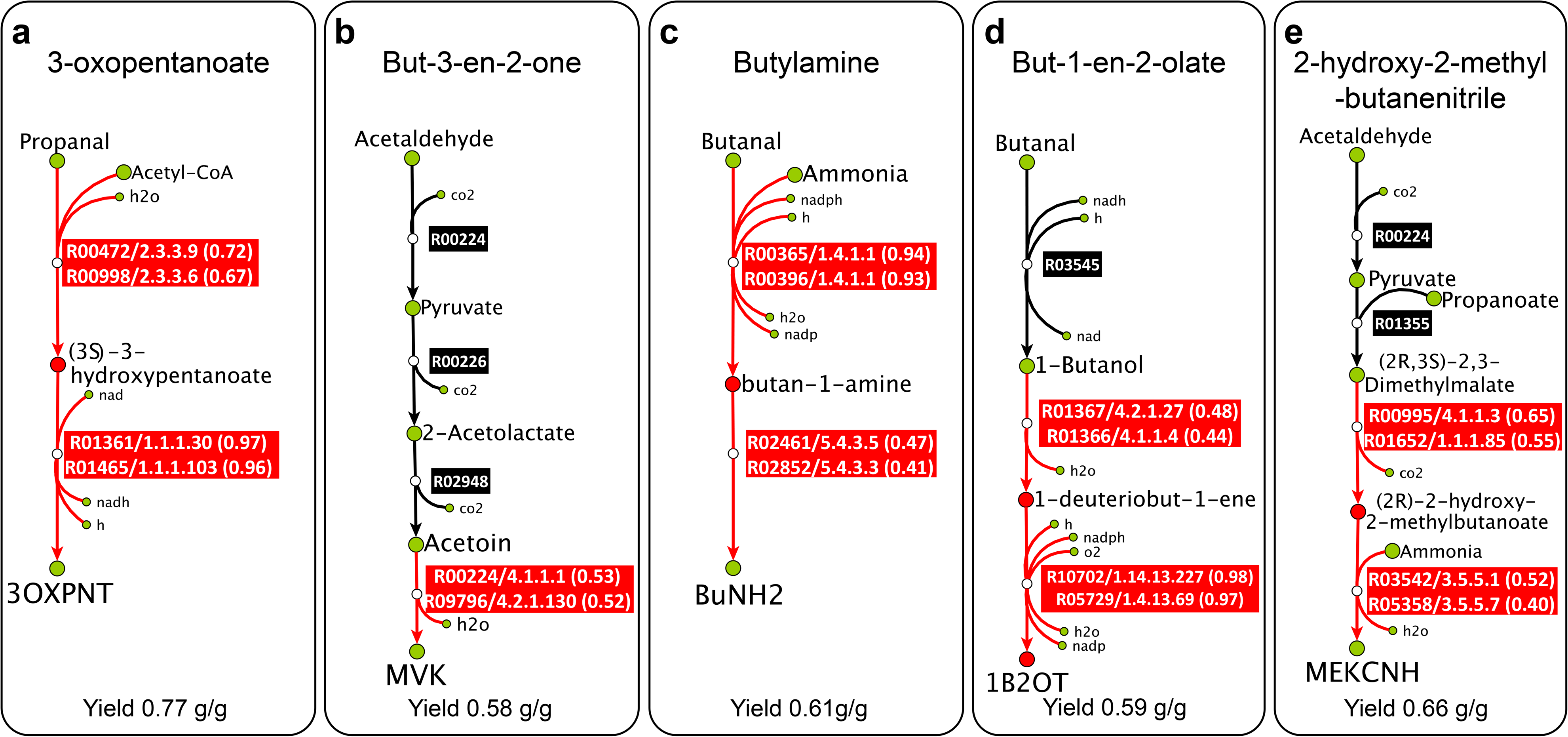
The highest ranked candidate pathways for the production of (**a**) 3OXPNT, (**b**) MVK, (**c**) BuNH2, (**d**) 1B2OT and (**e**) MEKCNH. Known non-orphan KEGG reactions are the black and novel reactions the red boxes. KEGG compounds are denoted as the green and PubChem compounds as the red circles. For each of the novel reactions, the KEGG IDs, catalyzing enzyme, and the BridgIT similarity scores of the two most similar non-orphan KEGG reactions are provided.

The pathways were visualized and can be consulted at http://lcsb-databases.epfl.ch/GraphList/ProjectList upon subscription.

### Further experimental implementation and pathway optimization

After ranking of the top candidate pathways, the experts can choose the most amenable ones for experimental implementation in the host organism. The implemented pathways typically need to be optimized further for economically viable production titers and rates. The optimization is performed through the Design-Built-Test-(Learn) cycle of metabolic engineering*^56–58^* where stoichiometric*^59–61^* and kinetic models*^62–69^*, genome editing*^70, 71^* and phenotypic characterization*^72^* are combined to improve recombinant strains for production of biochemicals.

## Methods

The BNICE.ch framework*^9, 10,20–25^* was employed to generate biosynthetic pathways towards 5 precursors of Methyl Ethyl Ketone: 3-oxopentanoate (3OXPNT), 2-hydroxy-2-methyl-butanenitrile (MEKCNH), but-3-en-2-one (MVK), 1-en-2-olate (1B2OT) and butylamine (BuNH_2_). We tested the set of reconstructed pathways against thermodynamic feasibility and mass balance constraints, and discarded the pathways that were not satisfying these requirements.*^9^* Next, the pruned pathways were ranked based on the several criteria, such as yield, number of known reaction steps and pathway length. The steps of the employed workflow are discussed further (Figure 6).

**Figure 6.**
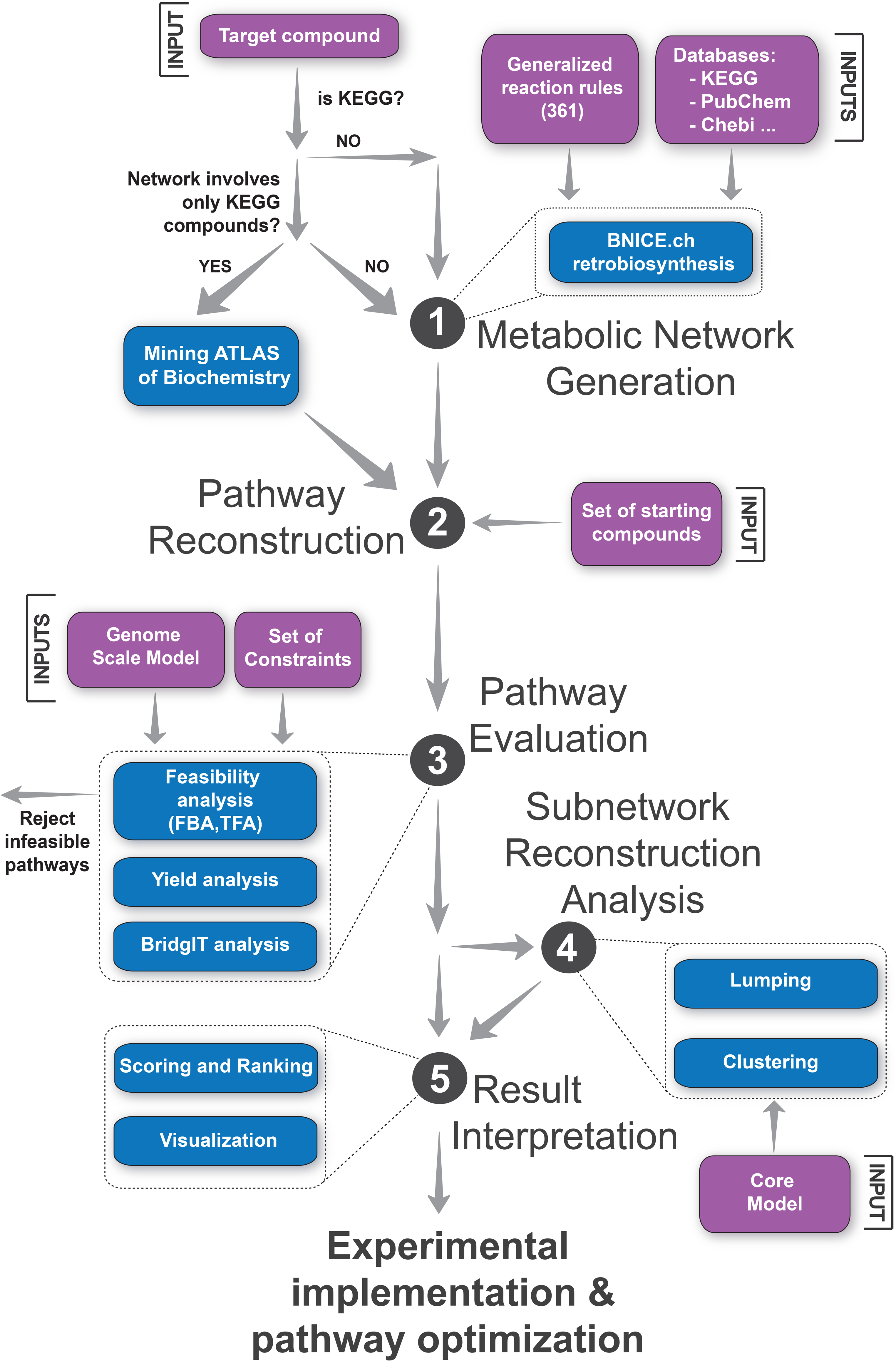
Computational pipeline for discovery, evaluation and analysis of biosynthetic pathways.

### Metabolic network generation

The retrobiosynthesis algorithm of BNICE.ch*^9, 48^* was applied to generate a biosynthetic network that contains all theoretically possible compounds and reactions that are up to 5 reaction steps away from MEK. The BNICE.ch network generation algorithm utilizes the expert-curated generalized enzyme reaction rules*^20, 21,73^* for identifying all potential compounds and reactions that lead to the production of the target molecules. The most recent version of BNICE.ch includes 361 bidirectional generalized reaction rules capable of reconstructing more than 6’500 KEGG reactions.*^23^* Starting from MEK and 26 cofactors required for the generalized enzyme reaction rules (Table S2 – Supporting Information), we identified the reactions that lead to MEK along with its potential precursors.^74^

Note that for studies where we need to generate a metabolic network that involves only KEGG compounds, mining the ATLAS of Biochemistry*^23^* is a more efficient procedure than using BNICE.ch retrobiosynthesis algorithm. The ATLAS of Biochemistry is a repository that contains all KEGG reactions and over 130’000 novel enzymatic reactions between KEGG compounds.

### Pathway reconstruction

The generated metabolic network was represented as a graph where the compounds were graph nodes and the reactions graph edges. A graph-based pathway search algorithm was then used to reconstruct all possible linear pathways up to the length of 4 reaction steps that connect the five target molecules with the set of 157 native *E. coli* metabolites (Table S3 – Supporting Information).*^41^* The search algorithm was employed as follows. Starting from a given native *E. coli* metabolite, we first searched for the shortest pathways toward a target molecule. Then, we searched for the pathways that are one reaction step longer than the shortest ones. We continued the search by gradually reconstructing the pathways of increased length, and we stopped with the 4 reaction step pathways. This procedure was repeated for all combinations of native *E. coli* metabolites and target molecules.

Note: If we were interested in pathways containing only KEGG reactions, we would perform a graph-based search over the network mined from the ATLAS of Biochemistry.^23^

### Pathway evaluation

It is crucial to identify and select, out of a vast number of generated pathways, the ones that satisfy physico-chemical constraints, such as mass balance and thermodynamics, or the ones that have an economically viable production yield of the target compounds from a carbon source. Evaluation of pathways is context-dependent, and it is important to perform it in an exact host organism model and under the same physiological conditions as the ones that will be used in the experimental implementation. Both Flux Balance Analysis (FBA)^42^ and Thermodynamic-based Flux Analysis (TFA)^43–47^ were performed to evaluate the pathways. We have also used BridgIT^50^ to identify candidate sequences for protein and evolutionary engineering in implementing the pathways. The availability of such sequences for the novel reactions and the ability to engineer them should also serve as a metric in ranking the feasibility of the pathways.

#### Flux balance and thermodynamic-based flux balance analysis

The generated pathways were embedded one at the time in the genome-scale model of *E. coli*, iJO1366*^41^* (File M1 – Supporting information) and FBA and TFA were performed for each the resulting models. In these analyses, we distinguished the following types the reactions: (*R1*) known and novel reactions for which have no information about their directionality; (*R2*) reactions that have preassigned directionality in iJO1366; and (R3) reactions that involve CO2 as a metabolite. It was assumed that the only carbon source was glucose and the following two types of constraints on reaction directionalities were applied:

> (*C1*) The preassigned reaction directionalities*^75^* from the iJO1366 model (*R2* reactions) were removed with the exception of ATP maintenance (ATPM), and it was assumed that the reactions that involve CO2 (*R3* reactions) are operating in the decarboxylation direction. The lower bound on ATPM was set to 8.39 mmol/gDCW/hr. The remaining reactions (*R1* reactions) were assumed to be bidirectional for FBA, whereas for TFA the directionality of these reactions was imposed by thermodynamics. The purpose of removing preassigned reaction directionalities was to investigate alternative hypotheses about the catalytic reversibility of the enzymes. The catalytic reversibility or irreversibility of enzymes could be altered through protein and evolutionary engineering and enzyme screening.*^45^* (C2) The preassigned directionalities of the *R2* reactions were kept and the directionality of the *R3* reactions was fixed towards decarboxylation.

Since FBA is computationally less expensive than TFA, we first performed FBA as a prescreening method to identify and discard the pathways: (i) that are not satisfying the mass balance, e.g., pathways that need co-substrates not present in the model; and (ii) that have a yield from glucose to the target compounds lower than a pre-specified threshold. In this work we used the pre-specified threshold of 0.1 mol/mol, however, this value can be chosen based on various criteria such as the economic viability of pathways. TFA was then performed on the reduced set of pathways to identify the pathways that are bio-energetically favorable and their yields from glucose to 5 target compounds were computed under thermodynamic constraints.

#### BridgIT analysis

We used BridgIT^50^ to associate genes to novel reactions appearing in the feasible pathways. BridgIT compares the similarity of a novel reaction to known reactions using the information about the structures of their substrates and products, and then assigns genes of the most similar known reactions as candidates for catalyzing the novel one. BridgIT integrates the information about the structures of substrates and products of a reaction into reaction difference fingerprints.^76^ These reaction fingerprints contain the information about chemical groups in substrates and products that were modified in the course of a reaction. BridgIT compares the reaction fingerprints of novel reactions to the ones of known reactions and quantifies this comparison with the Tanimoto similarity score. The Tanimoto score of 1 signifies that two compared reactions had a high similarity, whereas the Tanimoto score values close to 0 signify that there was no similarity. This score was used to rank the reactions identified as similar to each of the novel reactions. The gene and protein sequences of the highest ranked reactions were proposed as candidates for either a direct experimental implementation or enzyme engineering.

### Subnetwork reconstruction analysis

Once the biologically feasible pathways were identified and ranked, the parts of the metabolism that carry fluxes when the target compounds are produced from glucose were analyzed. We considered that the active parts of metabolism consisted of: (i) the core metabolic network (Figure 2a), which included the central carbon pathways, such as glycolysis, pentose phosphate pathway, tricarboxylic acid cycle, electron transport chain; and (ii) the active anabolic subnetworks (Figure 2a), which contain the reactions that would carry fluxes when a target molecule is produced but did not belong to the core metabolic network. We also defined the core precursors as metabolites that are connecting the core and the active anabolic subnetworks (Figure 2a).

The core metabolic network was derived from the genome-scale reconstruction iJO1366^41^ using the redGEM algorithm^77^, and the lumpGEM^53^ algorithm was then used to identify active anabolic subnetworks and to compute their lumped reactions. The analysis of lumped reactions allowed us to identify core precursors of the target chemicals. We then performed clustering to uncover core precursors, common enzymes, and intermediate metabolites of the anabolic subnetworks leading to the production of the target chemicals.

#### Identification and lumping of active anabolic subnetworks

The lumpGEM algorithm was applied to identify the comprehensive set of smallest metabolic subnetworks that were stoichiometrically balanced and capable of synthesizing a target compound from a defined set of core metabolites. The set of core metabolites belongs to the core metabolic network, and it includes also cofactors, small metabolites, and inorganic metabolites (Table S16 – Supporting Information). Then, for each target compound and for each identified subnetwork, we used lumpGEM to generate a corresponding lumped reaction. Within this process, the stoichiometric cost of core metabolites for the biosynthesis of these target compounds was also identified.

#### Clustering of subnetworks

To better understand the chemistry that leads towards the target compounds, we performed two types of clustering on the identified subnetworks:

- Clustering based on the structural similarity between the core precursors and byproducts of the lumped reactions. For each lumped reaction, we removed all non-carbon compounds, such as H2, O2, and phosphate, and the cofactor pairs, such as ATP and ADP, NAD+ and NADH, NADP+ and NADPH, flavodoxin oxidized and reduced, thioredoxin oxidized and reduced, ubiquinone and ubiquinol. This way, a set of substrates (core precursors) and byproducts of interest was created for each lumped reaction. We then used the *msim* algorithm from the RxnSim*^78^* tool to compare the lumped reactions based on individual similarities of their core precursors and byproducts. We finally used the obtained similarity scores to perform the clustering.
- Clustering based on the structural similarity between reactions that constitute the anabolic subnetworks. BridgIT was used to compute structural fingerprints of reactions that constitute the anabolic subnetworks, and we then performed a pairwise comparison of the anabolic subnetworks as follows. For a given pair of anabolic subnetworks, a pairwise comparison of their reactions was carried out. As a comparison metric we used the Tanimoto distance of the reaction fingerprints.*^79^* Based on this comparison, the pair of the most similar reactions in two subnetworks was found and the corresponding distance score was stored. This pair of reactions was then removed from comparison, and the next pair of the most similar reactions was found and their distance score was stored. We continued with this procedure until all pairs of reactions in two subnetworks were found. Whenever the number of reactions in two subnetworks was unequal, the unmatched reactions were ignored. The distance score between two compared subnetworks was formed as the sum of the distance scores of compared pairs of reactions. This procedure was repeated for all pairs of subnetworks. We then used the computed distance scores to perform the subnetworks clustering.

### Ranking and visualization of *in silico* pathways

In this step, we identified the pathways that were most likely to produce the target molecules. We defined the following criteria: (i) minimal number of reaction steps without promising candidate enzymes for catalyzing them; we consider that a novel reaction step has a promising candidate enzyme if the BridgIT algorithm has found its most similar known reactions with the similarity score higher than 0.3;*^50^* (ii) minimal number of novel reaction steps in a pathway; (iii) maximal yield from glucose to the target molecules; (iv) minimal number of reaction steps in the production pathway; and (v) highest average similarity scores of novel reaction steps from BridgIT. For scoring and ranking the biologically meaningful pathways, we used criterion (i) as the primary ranking. Then, equally ranked pathways from the primary ranking were further ranked based on criterion (ii). Analogously, we performed the tertiary, quaternary and quinary ranking based on criteria (iii), (iv) and (v), respectively.

An expert opinion is important in choosing the pathways for implementation. One can use other ranking criteria or a different prioritization of the criteria. For example, the pathways can be first ranked based on a maximum yield and then based on a minimal number of novel reactions. The complete set of pathways is provided on http://lcsb-databases.epfl.ch/GraphList/ProjectList, and the readers can rank the pathways according to their own criteria.

### Experimental implementation and pathway optimization

The highest ranked candidate pathways can then be experimentally implemented in the host organism and can further be optimized through the Design-Built-Test-(Learn) cycle of metabolic engineering.*^56–58^*

## Conclusions

In this work, we used BNICE.ch to reconstruct, evaluate and analyze more than 3.6 million biosynthetic pathways from the central carbon metabolites of *E. coli* towards five precursors of Methyl Ethyl Ketone (MEK), a 2^nd^ generation biofuel candidate. Our evaluation and analysis showed that more than 18’000 of these pathways are biologically feasible. We ranked these pathways based on process- and physiology-based criteria, and we identified gene and protein sequences of the structurally most similar KEGG reactions to the novel reactions in the feasible pathways, which can be used to accelerate their experimental realization. Implementation of the discovered pathways in *E. coli* will allow the sustainable and efficient production of five precursors of MEK (3OXPNT, MVK, 1B2OT, BuNH_2_, and MEKCNH), which can also be used as precursors for the production of other valuable chemicals.^36–35^

The pathway analysis methods developed and used in this work offer a systematic way for classifying and evaluating alternative ways for the production of target molecules. They also provide a better understanding of the underlying chemistry and can be used to guide the design of novel biosynthetic pathways for a wide range of biochemicals and for their implementation into host organisms.

The present study shows the potential of computational retrobiosynthesis tools for discovery and design of novel synthetic pathways for complex molecules, and their relevance for future developments in the area of metabolic engineering and synthetic biology.

## Author Information

### Author Contributions

V.H., L.M., N.H., M.A, and M.T. designed the study; M.T., N.H., and M.A. performed the experiments; V.H., L.M., N.H., M.A, M.T., L.M.B, B.E.E, and D.N. analyzed the data and wrote the manuscript.

### Notes

The authors declare no competing financial interests.

## Acknowledgments

We would like to thank Joana Pinto Vieira for her help with editing this manuscript, Homa MohammadiPeyhani for her help with generating reaction fingerprints for the clustering studies, and Jasmin Hafner for her help in publishing the BridgIT results on the web page.

## Supporting information

Tables S1-S16 (XLSX)

S1: List of biochemical and chemical compounds one step away from MEK.

S2: Initial set of compounds used for the generation of metabolic network around MEK.

S3: Stoichiometry of reactions that involve MEK as a metabolite.

S4: Reaction steps for a biochemical production of MEK from the five precursors (3OXPNT, MVK, BuNH_2_ 1B2OT, and MEKCNH) together with their most similar KEGG reactions.

S5: List of the starting compounds used in the pathway reconstruction.

S6: Yield histograms for 5 MEK precursors obtained with C1 constraints.

S7: Yield histograms for 5 MEK precursors obtained with C2 constraints.

S8: List of biomass building blocks in the genome-scale model of *E. coli* iJO1366.

S9: List of 185 alternative pathways from AcCoA and PpCoA to 3OXPNT.

S10: List of 115 pathways from acetate to 3OXPNT together with their classification in groups B1-B11.

S11. Top 5 ranked pathways toward 3OXPNT together with the BridgIT results for novel reaction steps.

S12. Top 5 ranked pathways toward MVK together with the BridgIT results for novel reaction steps.

S13. Top 5 ranked pathways toward BuNH_2_ together with the BridgIT results for novel reaction steps.

S14. Top 5 ranked pathways toward 1B2OT together with the BridgIT results for novel reaction steps.

S15. Top 5 ranked pathways toward MEKCNH together with the BridgIT results for novel reaction steps.

S16: List of metabolites in the core metabolic network.

Figures F1-F3 (PDF)

F1: Three alternative ways to produce 3OXPNT from acetate through 2 intermediate metabolites: 2-ethylmalate and 3-hydroxypentanoate.

F2: Three different pathways from acetate to 3OXPNT sharing the same lumped reaction.

F3: Clustering of all 242 alternatives for production of 3OXPNT from acetate.

File M1 (Matlab .mat file)

M1: Genome-scale model of *E. coli* iJO1366 and 242 active anabolic subnetworks connecting the core metabolism with 115 pathways from acetate to 3OXPNT together with their lumped reactions and stoichiometry.

## Abbreviations

| Abbreviation | Reaction | EC number |
| --- | --- | --- |
| 3OXCOAT | 3-oxoadipyl-CoA thiolase | 2.3.1.174 |
| ACACT1r | Acetyl-CoA C-acetyltransferase | 2.3.1.9 |
| ALDD3y | Aldehyde dehydrogenase (propanal, NADP) | 1.2.1.4 |
| AMPN | AMP nucleosidase | 3.2.2.4 |
| ASPK | Aspartate kinase | 2.7.2.4 |
| DADK | Deoxyadenylate kinase | 2.7.4.3 |
| DRPA | Deoxyribose-phosphate aldolase | 4.1.2.4 |
| FTHFLi | Formate-tetrahydrofolate ligase | 6.3.4.3 |
| LALDO3 | L-Lactaldehyde:NADP<sup>+</sup> 1-oxidoreductase | 1.1.1.283, 1.2.1.49 |
| MCITD | 2-methylcitrate dehydratase | 4.2.1.79 |
| MCITL2 | Methylisocitrate lyase | 4.1.3.30 |
| MGSA | Methylglyoxal synthase | 4.2.3.3 |
| MMM | Methylmalonyl-CoA mutase | 5.4.99.2 |
| NTD6 | 5'-nucleotidase (dAMP) | 3.1.3.89 |
| NTTP5 | Nucleoside triphosphate pyrophosphorylase | 3.6.1.19 |
| PPM2 | Phosphopentomutase 2 (deoxyribose) | 5.4.2.7 |
| PUNP2 | Purine-nucleoside phosphorylase | 2.4.2.1 |
| RNDR1 | Ribonucleoside-diphosphate reductase (ADP) | 1.17.4.1 |
| RNTR1c2 | Ribonucleoside-triphosphate reductase (ATP) | 1.17.4.2 |
| THRD_L | L-threonine deaminase | 4.1.1.19 |

| Abbreviation | Compound | Alternative name | Database IDs |  |
| --- | --- | --- | --- | --- |
|  |  |  | KEGG | PubChem |
| 3OXPNT | 3-Oxopentanoate | 3-Oxopentanoic acid | C02233 | 5297 |
| MVK | But-3-en-2-one | Methyl Vinyl Ketone | C20701 | 172232421 |
| BuNH<sub>2</sub> | Butylamine | Butanamine | C18706 | 124489380 |
| 1B2OT | But-1-en-2-olate | 1-Butene-2-olate | x | 54444500 |
| MEKCH | 2-Hydroxy-2-methylbutanenitrile | Methyl Ethyl Ketone Cyanohydrin | C18796 | 124489470 |
| 2dr1p | 2-deoxy-D-ribose-1-phosphate | 2-Deoxy-alpha-D-ribose 1-phosphate | C00672 | 3941 |
| 2dr5p | 2-deoxy-D-ribose 5-phosphate | 2-Deoxy-alpha-D-ribose 5-phosphate | C00673 | 3942 |
| dad | Deoxyadenosine | 2'-Deoxyadenosine | C00559 | 3839 |
| dgsn | Deoxyguanosine | 2'-Deoxyguanosine | C00330 | 3624 |
| duri | Deoxyuridine | 2'-Deoxyuridine | C00526 | 3809 |
| prpp | Phosphoribosyl pyrophosphate | 5-Phospho-alpha-D-ribose 1-diphosphate | C00119 | 3419 |
| ac | Acetate | Acetic acid | C00033 | 3335 |
| acCoA | Acetyl-CoA | Acetyl coenzyme A | C00024 | 3326 |
| akg | 2-oxoglutarate | 2-Ketoglutaric acid | C00026 | 3328 |
| asp-L | Aspartate | L-Aspartic acid | C00049 | 3351 |
| dhap | Dihydroxyacetone phosphate | 3-Hydroxy-2-oxopropyl phosphate | C00111 | 3411 |
| ppCoA | Propionyl-CoA | Propionyl coenzyme A | C00100 | 3400 |
| pyr | Pyruvate | 2-Oxopropanoate | C00022 | 3324 |
| r5p | Ribose-5-phosphate | alpha-D-Ribose 5-phosphate | C03736 | 6499 |
| succ | Succinate | Butanedionic acid | C00042 | 3344 |
| succCoA | Succinyl-CoA | Succinyl coenzyme A | C00091 | 3391 |

## References

1. Steen, E. J., Kang, Y., Bokinsky, G., Hu, Z., Schirmer, A., McClure, A., Del Cardayre, S. B., and Keasling, J. D. (2010) Microbial production of fatty-acid-derived fuels and chemicals from plant biomass, Nature 463, 559–562.

2. Lee, S. K., Chou, H., Ham, T. S., Lee, T. S., and Keasling, J. D. (2008) Metabolic engineering of microorganisms for biofuels production: from bugs to synthetic biology to fuels, Curr Opin Biotech 19, 556–563.

3. Yim, H., Haselbeck, R., Niu, W., Pujol-Baxley, C., Burgard, A., Boldt, J., Khandurina, J., Trawick, J. D., Osterhout, R. E., Stephen, R., Estadilla, J., Teisan, S., Schreyer, H. B., Andrae, S., Yang, T. H., Lee, S. Y., Burk, M. J., and Van Dien, S. (2011) Metabolic engineering of Escherichia coli for direct production of 1,4-butanediol, Nat Chem Biol 7, 445–452.

4. Atsumi, S., Hanai, T., and Liao, J. C. (2008) Non-fermentative pathways for synthesis of branched-chain higher alcohols as biofuels, Nature 451, 86–89.

5. Schirmer, A., Rude, M. a., Li, X., Popova, E., and del Cardayre, S. B. (2010) Microbial biosynthesis of alkanes, Science (New York, N.Y.) 329, 559–562.

6. Ghiaci, P., Norbeck, J., and Larsson, C. (2014) 2-Butanol and Butanone Production in *Saccharomyces cerevisiae* through Combination of a B12 Dependent Dehydratase and a Secondary Alcohol Dehydrogenase Using a TEV-Based Expression System, PLOS ONE 9, e102774.

7. Clomburg, J. M., and Gonzalez, R. (2010) Biofuel production in Escherichia coli: the role of metabolic engineering and synthetic biology, Appl Microbiol Biot 86, 419–434.

8. Cho, A., Yun, H., Park, J. H., Lee, S. Y., and Park, S. (2010) Prediction of novel synthetic pathways for the production of desired chemicals, Bmc Syst Biol 4.

9. Hadadi, N., and Hatzimanikatis, V. (2015) Design of computational retrobiosynthesis tools for the design of de novo synthetic pathways, Current Opinion in Chemical Biology 28, 99–104.

10. Henry, C. S., Broadbelt, L. J., and Hatzimanikatis, V. (2010) Discovery and Analysis of Novel Metabolic Pathways for the Biosynthesis of Industrial Chemicals: 3-Hydroxypropanoate, Biotechnol Bioeng 106, 462–473.

11. Moriya, Y., Shigemizu, D., Hattori, M., Tokimatsu, T., Kotera, M., Goto, S., and Kanehisa, M. (2010) PathPred: an enzyme-catalyzed metabolic pathway prediction server, Nucleic Acids Research 38, W138–W143.

12. Hou, B. K., Ellis, L. B. M., and Wackett, L. P. (2004) Encoding microbial metabolic logic: predicting biodegradation, Journal of Industrial Microbiology and Biotechnology 31, 261–272.

13. Ellis, L. B. M., Gao, J., Fenner, K., and Wackett, L. P. (2008) The University of Minnesota pathway prediction system: predicting metabolic logic, Nucleic Acids Research 36, W427–W432.

14. Campodonico, M. A., Andrews, B. A., Asenjo, J. A., Palsson, B. O., and Feist, A. M. (2014) Generation of an atlas for commodity chemical production in Escherichia coli and a novel pathway prediction algorithm, GEM-Path, Metabolic Engineering 25, 140–158.

15. Rodrigo, G., Carrera, J., Prather, K. J., and Jaramillo, A. (2008) DESHARKY: automatic design of metabolic pathways for optimal cell growth, Bioinformatics 24, 2554–2556.

16. Carbonell, P., Parutto, P., Herisson, J., Pandit, S. B., and Faulon, J. L. (2014) XTMS: pathway design in an eXTended metabolic space, Nucleic Acids Research 42, W389–W394.

17. Heath, A. P., Bennett, G. N., and Kavraki, L. E. (2010) Finding metabolic pathways using atom tracking, Bioinformatics 26, 1548–1555.

18. Dale, J. M., Popescu, L., and Karp, P. D. (2010) Machine learning methods for metabolic pathway prediction, Bmc Bioinformatics 11.

19. Prather, K. L. J., and Martin, C. H. (2008) De novo biosynthetic pathways: rational design of microbial chemical factories, Curr Opin Biotech 19, 468–474.

20. Hatzimanikatis, V., Li, C., Ionita, J. A., Henry, C. S., Jankowski, M. D., and Broadbelt, L. J. (2005) Exploring the diversity of complex metabolic networks, Bioinformatics 21, 1603–1609.

21. Hatzimanikatis, V., Li, C. H., Ionita, J. A., and Broadbelt, L. J. (2004) Metabolic networks: enzyme function and metabolite structure, Curr Opin Struc Biol 14, 300–306.

22. Hadadi, N., Soh, K. C., Seijo, M., Zisaki, A., Guan, X. L., Wenk, M. R., and Hatzimanikatis, V. (2014) A computational framework for integration of lipidomics data into metabolic pathways, Metabolic Engineering 23, 1–8.

23. Hadadi, N., Hafner, J., Shajkofci, A., Zisaki, A., and Hatzimanikatis, V. (2016) ATLAS of Biochemistry: A Repository of All Possible Biochemical Reactions for Synthetic Biology and Metabolic Engineering Studies, ACS Synthetic Biology, 1155–1166.

24. Soh, K. C., and Hatzimanikatis, V. (2010) Dreams of Metabolism, Trends Biotechnol 28, 501–508.

25. Brunk, E., Neri, M., Tavernelli, I., Hatzimanikatis, V., and Rothlisberger, U. (2012) Integrating computational methods to retrofit enzymes to synthetic pathways, Biotechnol Bioeng 109, 572–582.

26. Hoell, D., Mensing, T., Roggenbuck, R., Sakuth, M., Sperlich, E., Urban, T., Neier, W., and Strehlke, G. (2009) 2-Butanone, In Ullmann’sEncyclopedia of Industrial Chemistry, Wiley-VCH Verlag GmbH & Co. KGaA.

27. Hoppe, F., Burke, U., Thewes, M., Heufer, A., Kremer, F., and Pischinger, S. (2016) Tailor-Made Fuels from Biomass: Potentials of 2-butanone and 2-methylfuran in direct injection spark ignition engines, Fuel 167, 106–117.

28. Srirangan, K., Liu, X., Akawi, L., Bruder, M., Moo-young, M., and Chou, C. P. (2016) Engineering Escherichia coli for Microbial Production of Butanone, 2574–2584.

29. Yoneda, H., Tantillo, D. J., and Atsumi, S. (2014) Biological production of 2-butanone in Escherichia coli, ChemSusChem 7, 92–95.

30. Multer, A., McGraw, N., Hohn, K., and Vadlani, P. (2013) Production of Methyl Ethyl Ketone from Biomass Using a Hybrid Biochemical/Catalytic Approach, Industrial & Engineering Chemistry Research 52, 56–60.

31. Drabo, P., Tiso, T., Heyman, B., Sarikaya, E., Gaspar, P., Förster, J., Büchs, J., Blank, L. M., and Delidovich, I. (2017) Anionic Extraction for Efficient Recovery of Biobased 2,3-Butanediol—A Platform for Bulk and Fine Chemicals, ChemSusChem 10, 3252–3259.

32. Kanehisa, M., Sato, Y., Kawashima, M., Furumichi, M., and Tanabe, M. (2016) KEGG as a reference resource for gene and protein annotation, Nucleic Acids Research 44, D457–D462.

33. Kanehisa, M., and Goto, S. (2000) KEGG: Kyoto Encyclopedia of Genes and Genomes, Nucleic Acids Research 28, 27–30.

34. Engel, T. (2007) The structural- and bioassay database PubChem, Nachr Chem 55, 521–524.

35. Kim, S., Thiessen, P. A., Bolton, E. E., Chen, J., Fu, G., Gindulyte, A., Han, L., He, J., He, S., Shoemaker, B. A., Wang, J., Yu, B., Zhang, J., and Bryant, S. H. (2016) PubChem Substance and Compound databases, Nucleic Acids Research 44, D1202–D1213.

36. Krumpfer, J. W., Giebel, E., Frank, E., Müller, A., Ackermann, L.-M., Tironi, C. N., Mourgas, G., Unold, J., Klapper, M., Buchmeiser, M. R., and Müllen, K. (2017) Poly(Methyl Vinyl Ketone) as a Potential Carbon Fiber Precursor, Chemistry of Materials 29, 780–788.

37. Siegel, H., and Eggersdorfer, M. (2000) Ketones, In Ullmann’sEncyclopedia of Industrial Chemistry, Wiley-VCH Verlag GmbH & Co. KGaA.

38. Eller, K., Henkes, E., Rossbacher, R., and Höke, H. (2000) Amines, Aliphatic, In Ullmann’s Encyclopedia of Industrial Chemistry, Wiley-VCH Verlag GmbH & Co. KGaA.

39. Chaparro-Riggers, J. F., Rogers, T. A., Vazquez-Figueroa, E., Polizzi, K. M., and Bommarius, A. S. (2007) Comparison of three enoate reductases and their potential use for biotransformations, Adv Synth Catal 349, 1521–1531.

40. Lilley, D. M. J., Clegg, R. M., Diekmann, S., Seeman, N. C., Von Kitzing, E., and Hagerman, P. J. (1995) A nomenclature of junctions and branchpoints in nucleic acids, Nucleic Acids Research 23, 3363–3364.

41. Orth, J. D., Conrad, T. M., Na, J., Lerman, J. A., Nam, H., Feist, A. M., and Palsson, B. O. (2011) A comprehensive genome-scale reconstruction of Escherichia coli metabolism-2011, Mol Syst Biol 7.

42. Orth, J. D., Thiele, I., and Palsson, B. Ø. (2010) What is flux balance analysis?, Nature biotechnology 28, 245–248.

43. Henry, C. S., Broadbelt, L. J., and Hatzimanikatis, V. (2007) Thermodynamics-based metabolic flux analysis, Biophys J 92, 1792–1805.

44. Henry, C. S., Jankowski, M. D., Broadbelt, L. J., and Hatzimanikatis, V. (2006) Genome-scale thermodynamic analysis of Escherichia coli metabolism, Biophys J 90, 1453–1461.

45. Ataman, M., and Hatzimanikatis, V. (2015) Heading in the right direction: thermodynamics-based network analysis and pathway engineering, Curr Opin Biotech 36, 176–182.

46. Soh, K. C., and Hatzimanikatis, V. (2010) Network thermodynamics in the post-genomic era, Curr Opin Microbiol 13, 350–357.

47. Soh, K. S., and Hatzimanikatis, V. (2014) Constraining the flux space using thermodynamics and integration of metabolomics data, Methods in Molecular Biology 1191, 49–63.

48. Islam, M. A., Hadadi, N., Ataman, M., Hatzimanikatis, V., and Stephanopoulos, G. (2017) Exploring biochemical pathways for mono-ethylene glycol (MEG) synthesis from synthesis gas, Metabolic Engineering 41, 173–181.

49. Demirel, Y. s. (2014) Nonequilibrium thermodynamics transport and rate processes in physical, chemical and biological systems, 3rd ed., Elsevier, Amsterdam.

50. Hadadi, N., MohamadiPeyhani, H., Miskovic, L., Seijo, M., and Hatzimanikatis, V. (2017) Knowledge of the Neighborhood of the Reactive Site up to Three Atoms Can Predict Biochemistry and Protein Sequences, bioRxiv, https://doi.org/10.1101/210039.

51. Neidhardt, F. C., Ingraham, J. L., and Schaechter, M. (1990) Physiology of the bacterial cell: a molecular approach, Sinauer Associates, Sunderland, Mass.

52. Sudarsan, S., Dethlefsen, S., Blank, L. M., Siemann-Herzberg, M., and Schmid, A. (2014) The Functional Structure of Central Carbon Metabolism in Pseudomonas putida KT2440, Appl Environ Microb 80, 5292–5303.

53. Ataman, M., and Hatzimanikatis, V. (2017) lumpGEM: Systematic generation of subnetworks and elementally balanced lumped reactions for the biosynthesis of target metabolites, Plos Comput Biol 13, e1005513.

54. Chen, Y., Daviet, L., Schalk, M., Siewers, V., and Nielsen, J. (2013) Establishing a platform cell factory through engineering of yeast acetyl-CoA metabolism, Metab Eng 15, 48–54.

55. Haller, T., Buckel, T., Rétey, J., and Gerlt, J. A. (2000) Discovering New Enzymes and Metabolic Pathways: Conversion of Succinate to Propionate by Escherichia coli, Biochemistry 39, 4622–4629.

56. Petzold, C. J., Chan, L. J. G., Nhan, M., and Adams, P. D. (2015) Analytics for Metabolic Engineering, Frontiers in Bioengineering and Biotechnology 3, 135.

57. Campbell, K., Xia, J., and Nielsen, J. The Impact of Systems Biology on Bioprocessing, Trends Biotechnol 35, 1156–1168.

58. Miskovic, L., Alff-Tuomala, S., Soh, K. C., Barth, D., Salusjarvi, L., Pitkanen, J. P., Ruohonen, L., Penttila, M., and Hatzimanikatis, V. (2017) A design-build-test cycle using modeling and experiments reveals interdependencies between upper glycolysis and xylose uptake in recombinant S. cerevisiae and improves predictive capabilities of large-scale kinetic models, Biotechnol Biofuels 10.

59. Bordbar, A., Monk, J. M., King, Z. A., and Palsson, B. O. (2014) Constraint-based models predict metabolic and associated cellular functions, Nat Rev Genet 15, 107–120.

60. Maarleveld, T. R., Khandelwal, R. A., Olivier, B. G., Teusink, B., and Bruggeman, F. J. (2013) Basic concepts and principles of stoichiometric modeling of metabolic networks, Biotechnol J 8, 997–U952.

61. Garcia-Albornoz, M. A., and Nielsen, J. (2013) Application of Genome-Scale Metabolic Models in Metabolic Engineering, Industrial Biotechnology 9, 203–214.

62. Miskovic, L., Tokic, M., Fengos, G., and Hatzimanikatis, V. (2015) Rites of passage: requirements and standards for building kinetic models of metabolic phenotypes, Curr Opin Biotech 36, 146–153.

63. Miskovic, L., and Hatzimanikatis, V. (2010) Production of biofuels and biochemicals: in need of an ORACLE, Trends Biotechnol 28, 391–397.

64. Andreozzi, S., Chakrabarti, A., Soh, K. C., Burgard, A., Yang, T. H., Van Dien, S., Miskovic, L., and Hatzimanikatis, V. (2016) Identification of metabolic engineering targets for the enhancement of 1,4-butanediol production in recombinant E. coli using large-scale kinetic models, Metabolic Engineering 35, 148–159.

65. Chakrabarti, A., Miskovic, L., Soh, K. C., and Hatzimanikatis, V. (2013) Towards kinetic modeling of genome-scale metabolic networks without sacrificing stoichiometric, thermodynamic and physiological constraints, Biotechnol J 8, 1043–1057.

66. Savoglidis, G., dos Santos, A. X. D., Riezman, I., Angelino, P., Riezman, H., and Hatzimanikatis, V. (2016) A method for analysis and design of metabolism using metabolomics data and kinetic models: Application on lipidomics using a novel kinetic model of sphingolipid metabolism, Metabolic Engineering 37, 46–62.

67. Stanford, N. J., Lubitz, T., Smallbone, K., Klipp, E., Mendes, P., and Liebermeister, W. (2013) Systematic Construction of Kinetic Models from Genome-Scale Metabolic Networks, PLOS One 8.

68. Dash, S., Khodayari, A., Zhou, J., Holwerda, E. K., Olson, D. G., Lynd, L. R., and Maranas, C. D. (2017) Development of a core Clostridium thermocellum kinetic metabolic model consistent with multiple genetic perturbations, Biotechnol Biofuels 10.

69. Lee, Y., Lafontaine Rivera, J. G., and Liao, J. C. (2014) Ensemble Modeling for Robustness Analysis in engineering non-native metabolic pathways, Metabolic Engineering 25, 63–71.

70. Esvelt, K. M., and Wang, H. H. (2013) Genome-scale engineering for systems and synthetic biology, Mol Syst Biol 9.

71. Barrangou, R., and Doudna, J. A. (2016) Applications of CRISPR technologies in research and beyond, Nature Biotechnology 34, 933–941.

72. Bochner, B. R. (2009) Global phenotypic characterization of bacteria, Fems Microbiol Rev 33, 191–205.

73. Li, C. H., Henry, C. S., Jankowski, M. D., Ionita, J. A., Hatzimanikatis, V., and Broadbelt, L. J. (2004) Computational discovery of biochemical routes to specialty chemicals, Chem Eng Sci 59, 5051–5060.

74. Corey, E. J. (1991) The Logic of Chemical Synthesis – Multistep Synthesis of Complex Carbogenic Molecules, Angew Chem Int Edit 30, 455–465.

75. Frainay, C., and Jourdan, F. (2017) Computational methods to identify metabolic sub-networks based on metabolomic profiles, Briefings in Bioinformatics 18, 43–56.

76. James, C. A., and Weininger, D. Daylight Theory Manual, Daylight Chemical Information Systems, Inc.: Irvine, CA.

77. Ataman, M., Gardiol, D. H. F., Fengos, G., and Hatzimanikatis, V. (2017) redGEM: Systematic Reduction and Analysis of Genome-scale Metabolic Reconstructions for Development of Consistent Core Metabolic Models, Plos Comput Biol, e1005444.

78. Giri, V., Sivakumar, T. V., Cho, K. M., Kim, T. Y., and Bhaduri, A. (2015) RxnSim: a tool to compare biochemical reactions, Bioinformatics 31, 3712–3714.

79. Bajusz, D., Rácz, A., and Héberger, K. (2015) Why is Tanimoto index an appropriate choice for fingerprint-based similarity calculations?, Journal of Cheminformatics 7, 20.

